# INHBA/Activin A promotes tumor growth and induces resistance to anti-PD-L1 therapy by suppressing IFN-γ signaling

**DOI:** 10.1101/2023.12.07.570561

**Authors:** Fanglin Li, Longhua Gu, Yongliang Tong, Xiaolu Yu, Runqiu Chen, Nan Liu, Shiyi Chen, Jiangling Lu, Yuan Si, Jianhua Sun, Jing Chen, Yiru Long, Likun Gong

## Abstract

Inhibin beta A (INHBA) and its homodimer activin A have pleiotropic effects on modulation of immune responses and tumor progression, respectively, but it remains uncertain whether tumors may release activin A to regulate anti-tumor immunity. As evidenced by our RNA-Seq and in vitro results, the interferon-γ (IFN-γ) signaling pathway was significantly down-regulated by tumor intrinsic activin A. Tumor INHBA deficiency led to lower expression of PD-L1 induced by IFN-γ, resulting in poor responsiveness to anti-PD-L1 therapy. On the other hand, decreased secretion of IFN-γ-stimulated chemokines, including C-X-C motif chemokine 9 (CXCL9) and 10 (CXCL10), impaired the infiltration of effector T cells into the tumor microenvironment. Furthermore, the activin A-specific antibody garetosmab improved anti-tumor immunity and its combination with the anti-PD-L1 antibody atezolizumab showed a superior therapeutic effect to monotherapy. Our findings reveal that INHBA/activin A is involved in anti-tumor immunity by inhibiting the IFN-γ signaling pathway and considered to be a potential target to overcome anti-PD-L1 resistance in clinical cancer treatment.

## Introduction

Immunotherapy based on programmed death ligand 1 (PD-L1) blockade has made dramatic breakthroughs in multiple cancer types, including non-small cell lung cancer (NSCLC), urothelial carcinoma and triple-negative breast cancer (TNBC) ^1, 2, 3^. However, a majority of solid tumor patients experience poor responsiveness to anti-PD-L1 therapy or fail to obtain durable therapeutic effect ^4^. In recent years, several factors accounting for clinical resistance to treatment targeting the PD-1/PD-L1 axis have been listed, such as weak tumor immunogenicity, the PD-1/PD-L1 independent T cell evasion mechanism and low tumor intrinsic PD-L1 expression ^5, 6, 7^. Focusing on these factors, concerted efforts have been put into the uncovering of novel targets and the development of optimized combination strategies to address the difficulties encountered in PD-L1 blockage.

Activin A, which is encoded by *Inhba*, is a homodimer of the inhibin beta A subunit and functions as a critical member of transforming growth factor-β (TGF-β) superfamily ^8^. Activin A switches on its signal by binding to the receptor complex composed of activin receptor-like kinase 4 (ALK4), activin receptor type IIB (ActRIIB) and activin receptor type IIA (ActRIIA).

Subsequently, it activates phosphorylation of mothers against decapentaplegic (SMAD) 2 and 3 to initiate transcription of genes including *Pax6*, *FST* and *p21* ^9, 10, 11, 12^. Originally, activin A was seen as a gonadal hormone that could modulate the secretion of follicle-stimulating hormone (FSH) ^13^.

Increasing evidence has revealed the involvement of activin A in immune regulation and tumor progression. As to innate immunity, activation of macrophages or dendritic cells (DCs) can boost the release of activin A, which in turn suppresses phagocytosis and the antigen-presenting ability of macrophages ^14, 15, 16^. Activin A can also skew macrophage differentiation toward the M2 type ^17^. Surprisingly, activin A-treated DC vaccine inhibits tumor growth ^18^. For adaptive immunity, activin A dampens the activity and function of CD8^+^ T cells ^19^. Moreover, it serves as a Th2 cytokine and can regulate the differentiation of pathogenic Th17 ^20, 21^. Contrary to TGF-β, activin A tends to induce upregulation of IL-10-producing type 1 regulatory T (Tr1) cells instead of natural regulatory T cells (nTregs) in order to exert immunosuppressive function^22^. Clinically, *INHBA* expression is significantly correlated with the reduced overall survival rate, poor prognosis and low immune infiltrating level of patients suffering from breast cancer, cervical cancer and colon adenocarcinoma (COAD) ^23, 24, 25^. But in-depth study of whether activin A/INHBA affects tumor growth via immune-related pathways still lacks. Clinical bioinformatics analysis has demonstrated that *INHBA* is upregulated in melanoma patients who are resistant to immune checkpoint blockade therapy (ICB) ^26^. A recent study also pointed out that activin A could impair immune therapy response in a CD8^+^ T cell-dependent way in a melanoma model ^27^. However, concrete mechanisms and expanding evidence on other tumor types needs further exploration.

In our research, we investigated the effects and mechanisms of tumor intrinsic INHBA on carcinogenesis, tumor immunity and anti-PD-L1 therapy, hoping to inspire new insights into how INHBA regulates tumor immunity and to assess the value of INHBA-targeting strategies in improving the responsive rate of PD-1/PD-L1 blockade.

## Results

### INHBA promotes tumor growth via T cell-related pathways

To determine whether *Inhba* expression in tumor tissues is related to the malignant progression and poor prognosis of cancer patients, we conducted a comparative analysis of INHBA expression levels in tumor tissues and normal tissues across 33 different cancer types by GEPIA tool^28^ based on the TCGA database. The results showed that *INHBA* is elevated in multiple tumor types, including breast cancer (BRCA) and colon adenocarcinoma (COAD) (Fig1A). In addition, survival analysis also corroborates that *INHBA* expression is negatively correlated with the prognosis of many types of cancer patients (FigS1A).

**Fig. 1.**
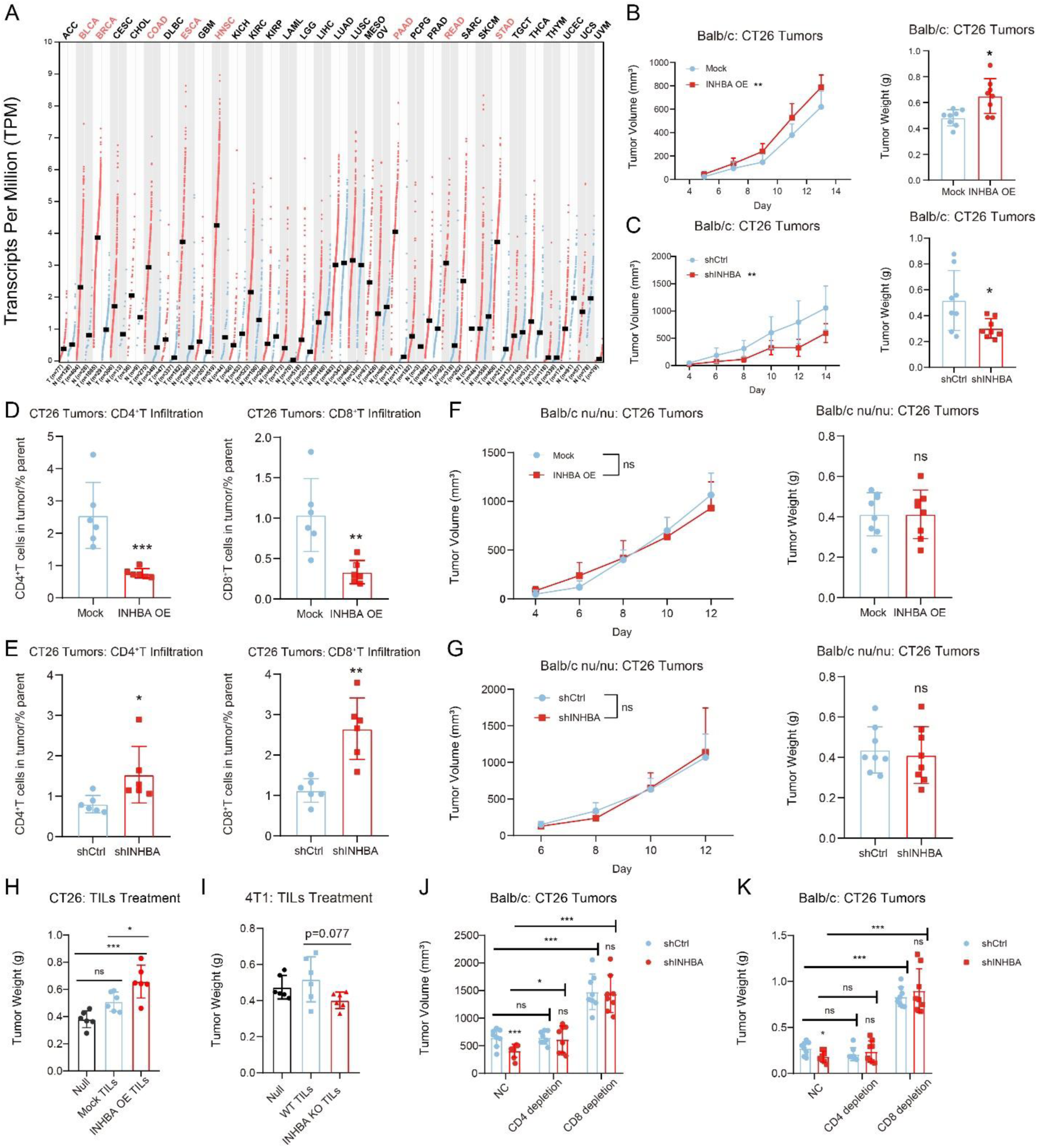
INHBA promoted tumor growth via T cell-related immunity. (**A**) Human *INHBA* mRNA levels in different tumor types from the TCGA database were compared to normal tissue *INHBA* expression and were determined by GEPIA. The red font shows the tumors with a statistically significant difference in comparison by the Wilcoxon test. (**B**) Tumor growth kinetics (left panel) and weights (right panel) of CT26 mock cells versus CT26 INHBA OE cells in Balb/c mice (n = 8). (**C**) Tumor growth kinetics (left panel) and weights (right panel) of CT26 cells expressing non-targeting shRNA (CT26 shCtrl) versus CT26 cells expressing INHBA shRNA (CT26 shINHBA) in Balb/c mice (n = 8). (**D** to **E**) Quantification of CD4^+^ T cells and CD8^+^ T cells infiltration in mock CT26 and INHBA OE CT26 tumors (D) or shCtrl CT26 and shINHBA CT26 tumors (E). (**F**) Tumor growth kinetics (left panel) and weights (right panel) of mock CT26 tumors and INHBA OE CT26 tumors in Balb/c nu/nu mice (n = 8). (**G**) Tumor growth kinetics (left panel) and weights (right panel) of shCtrl CT26 tumors and shINHBA CT26 tumors in Balb/c nu/nu mice (n = 8). (**H** to **I**) The weight of CT26 tumors (H) or 4T1 tumors (I) in Balb/c nu/nu mice injected with adoptive TILs are shown (n = 6). TILs isolated from mock or INHBA OE tumors were peritumorally injected into respective tumors. (**J** to **K**) Mice bearing shCtrl or shINHBA CT26 tumors were dosed with CD4^+^ T or CD8^+^ T cells depletion reagents. Tumor volume (J) and tumor weight (K) are shown (n = 8). The data are presented as the mean ± SEM. * p < 0.05; ** p < 0.01; *** p < 0.001; ns not significant by unpaired t test or one-way/two-way ANOVA followed by the Tukey multiple comparisons test.

We constructed several mouse *Inhba* gene-editing tumor cell lines in order to study the effect of INHBA on tumor progression (FigS1B). In the cell proliferation assay, in vitro growth of CT26 was not altered by *Inhba* overexpression (OE) or knockdown (FigS1C). By subcutaneously transplanting CT26 into BALB/c mice, we found that *Inhba* overexpression resulted in a stark increase in tumor size and weight, while *Inhba* knockdown suppressed tumor growth with a 41.66% inhibition rate (Fig1B and 1C). The same results were further confirmed in the B16 and MC38 models (FigS1D).

By employing the TIMER 2.0 tool^29^, we figured out that in most cancer types, *Inhba* expression in tumors was associated inversely with the infiltration of CD4^+^ T and CD8^+^ T cells (FigS2A). In accordance with the clinical data, immunophenotyping by flow cytometry uncovered a negative correlation between intratumoral *Inhba* expression and CD4^+^ T and CD8^+^ T cell infiltration (Fig1D, Fig1E and FigS3A-S3B). Next, we inoculated CT26 *Inhba* gene-editing cells into T cell-deficient BALB/c nu/nu mice and found that the influences imparted by INHBA on tumor growth were ablated (Fig1F and 1G). Considering activin A coded by *Inhba* is a typical Th2 cytokine and can induce differentiation of immunosuppressive Tr1 cells^20, 22^, we sought to determine whether INHBA could also dampen T cell function. Consequently, the overexpression of *Inhba* effectively promoted Th2 differentiation and Tr1 infiltration, and it also inhibited the cytotoxicity of CD8^+^ T cells (FigS3C-S3E). We also found that tumor *Inhba* overexpression brought about a higher splenic coefficient (FigS3F). And as anticipated, the serum concentration of INHBA is markedly increased in the *Inhba* overexpression group (FigS3G). Thus, given that activin A is a secretory protein, we speculated that it would be secreted into the periphery and alter systemic immunity.

Adoptive transfer of tumor-infiltrating lymphocytes (TILs) into wild-type (WT) CT26 bearing BALB/c nu/nu further underpinned the suppressive role of INHBA on T cell function (Fig1H and 1J). Depletion of CD4^+^ T and CD8^+^ T cells highlighted the pivotal roles of these two cell types in the involvement of INHBA-induced tumor progression (Fig1K, 1L and FigS4). These data indicate that INHBA regulates tumor growth in a T cell dependent way.

### INHBA induces resistance to anti-PD-L1 therapy

Analysis of clinical cohorts conducted by Riaz N et al.^30^ and Hugo W et al.^5^ revealed that INHBA was observably upregulated in tumor tissues of melanoma patients non-responsive to ICB therapy (Fig2A and FigS5A), which led us to make the assumption that INHBA may mediate resistance to immune checkpoint blockade (ICB) therapy.

**Fig. 2.**
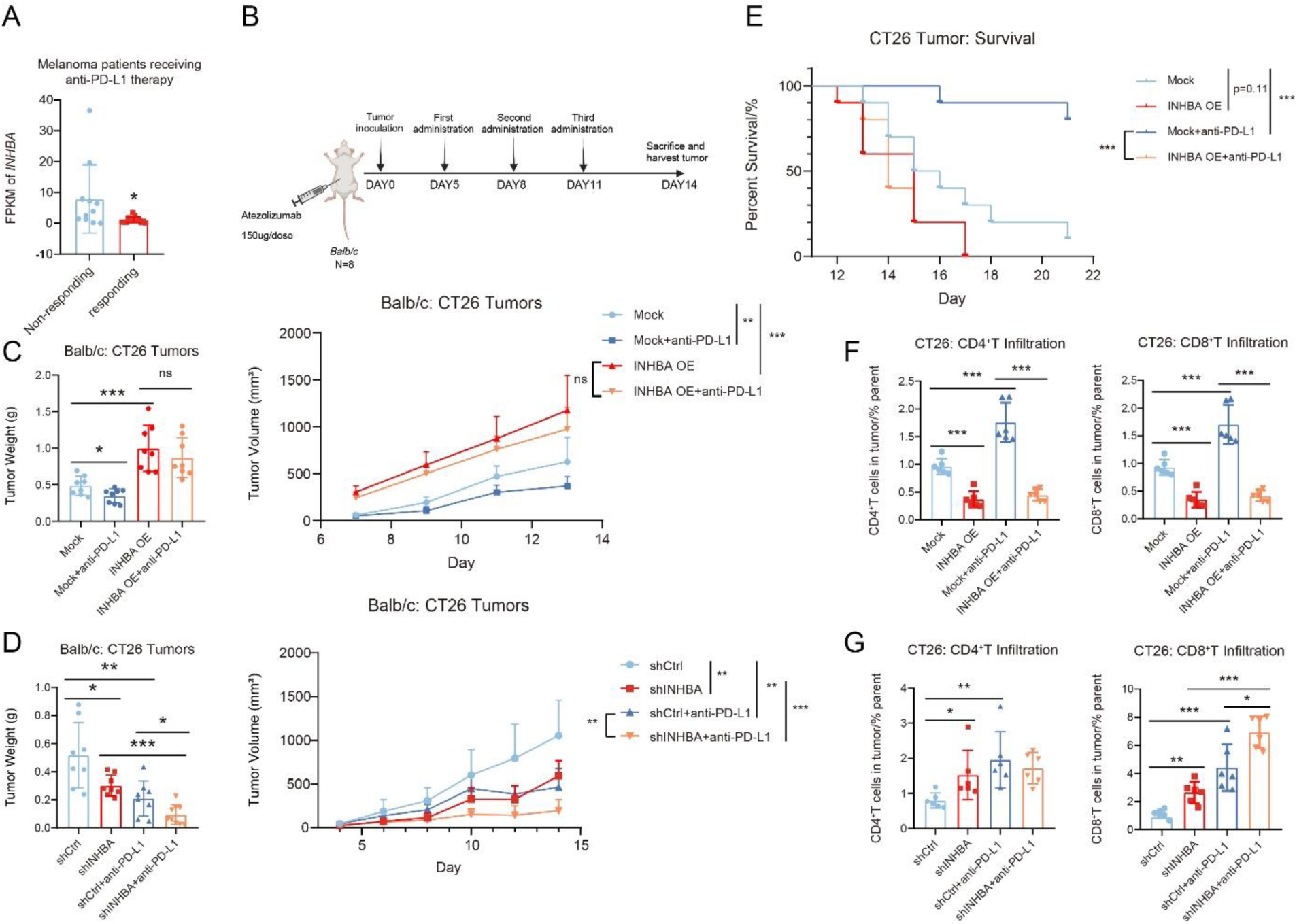
INHBA induced resistance to anti-PD-L1 therapy. (**A**) INHBA mRNA expression was compared between tumor from melanoma patients responsive or non-responsive to anti-PD-1/PD-L1 therapy provided by available GEO data (GSE78220). (**B**) Schematic diagram of administration of atezolizumab. Atezolizumab was intraperitoneal injected every 3 day to tumor-bearing mice (n = 8). (**C**) Tumor weights (left panel) and growth kinetics (right panel) of mock or INHBA OE CT26 tumors with treatment of atezolizumab or not (n = 8). (**D**) Tumor weights (left panel) and growth kinetics (right panel) of shCtrl or shINHBA CT26 tumors with treatment of atezolizumab or not (n = 8). (**E**) Kaplan–Meier plot of survival is shown for mock or INHBA OE CT26 tumors with or without treatment of atezolizumab (n = 10). (**F-G**) Quantification of CD4^+^T cells and CD8^+^T cells infiltration from mock or INHBA OE CT26 tumors (F) and shCtrl or shINHBA CT26 tumors (G) with or without treatment of atezolizumab (n = 6). The data are presented as the mean ±SEM. * p < 0.05; ** p < 0.01; *** p < 0.001; ns not significant by unpaired t test or one-way ANOVA followed by Tukey’s multiple comparisons test. P values of survival plots were calculated by the log-rank test.

To verify the above hypothesis, we treated tumor-bearing mice with the PD-L1 antibody atezolizumab (Fig2B). The results demonstrated that overexpression of tumor INHBA eliminated the anti-tumor function presented by atezolizumab, while INHBA deficiency enhanced the efficacy of anti-PD-L1 therapy (Fig2C, Fig2D, FigS5B, and S5C), which is consistent with our previous supposition. Survival analysis further consolidated this idea (Fig2E). As we assumed that the tumor-derived INHBA effect on tumor progression is contingent upon T cell-mediated immunity, flow cytometry was used to test if INHBA-induced anti-PD-L1 resistance was also associated with immunosuppression of T cells. The atezolizumab-induced improvement in T cell infiltration was evidently nullified by tumor INHBA. Instead, the knockdown of INHBA enhanced especially CD8^+^ T cell infiltration after anti-PD-L1 administration, with a minor impact on CD4^+^ T cells (Fig2F, Fig2G, FigS5D, and S5E). In terms of modulation of T cell function, we further confirmed that INHBA augmented Tr1 infiltration, although PD-L1 blockade slightly inhibited this effect (FigS5F). Taken together, tumor INHBA contributes to anti-PD-L1 resistance.

### INHBA inhibits IFN-γ response pathway

To explore the mechanism of how INHBA is involved in tumor progression and anti-PD-L1 resistance, we conducted bulk RNA sequencing (RNA-Seq) on tumor tissues and TILs, respectively, to probe the key signaling pathway. We observed that a variety of immune-related pathways were downregulated in the *Inhba* overexpression group, including antigen processing, chemokines and interferon (Fig3A and FigS6A). GSEA enrichment analysis showed that no matter whether treated with PD-L1 antibody or not, *Inhba* overexpression conspicuously suppressed IFN-γ response pathway (Fig3B). For both tumors and TILs, most of the genes encompassed in IFN-γ response pathway exhibit lower expression in *Inhba* overexpression group in comparison to the control (Fig3C and FigS6B). As we expected, anti-PD-L1 therapy markedly upregulated many immune pathways in the mock group; however, it conferred no alternation in *Inhba* overexpression context, which again indicated that INHBA led to the unresponsiveness to anti-PD-L1 treatment (Fig3A). Consistently, the immune score of IFN-γ signaling pathway reflected the same trend (Fig 3D and Fig S6C). These results collectively prompted us to speculate that INHBA triggers tumor progression and anti-PD-L1 resistance via IFN-γ response pathway.

**Fig. 3.**
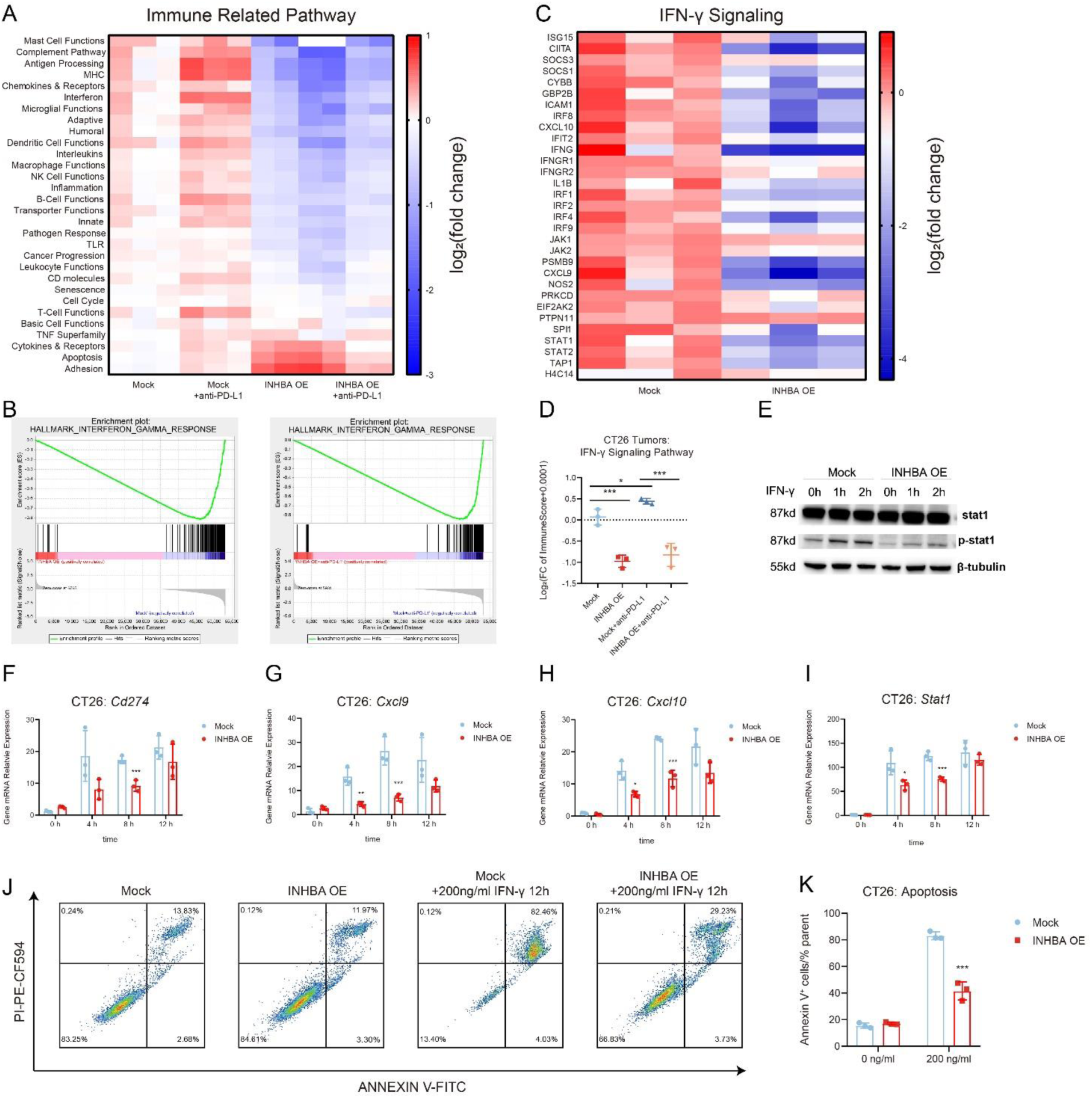
INHBA inhibited IFN-γ response pathway. (**A**) Signature scores [defined as the mean log_2_(fold change) among all genes measured by RNA-seq in the signature] for immune-associated processes are shown as heat map (n = 3). Samples are from tumors of CT26 bearing mice. (**B**) GSEA analysis shows decreased IFN-γ signaling in INHBA OE CT26 tumors compared with mock CT26 tumors. (**C**) Signature scores for WP_TYPE_II_INTERFERON_SIGNALING_IFNG geneset are shown as scatter plot (n = 3). Samples are from tumors of CT26 bearing mice. (**D**) Expression scores [defined as log_2_(fold change) of all genes measured by RNA-seq in the signature] for all genes in WP_TYPE_II_INTERFERON_SIGNALING_IFNG geneset are shown as heat map (n = 3). Samples are from tumors of CT26 bearing mice. (**E**) Western blot analysis of p-STAT1 and STAT1 expression in mock or INHBA OE CT26 cells with treatment of IFN-γ. β-Tubulin was used as the protein loading control. (**F** to **I**) Gene expression for *Cd274* (F), *Cxcl9* (G), *Cxcl10* (H) and *Stat1* (I) in mock or INHBA OE CT26 cells with IFN-γ treatment (n = 3). (**J** to **K**) Representative histograms (J) and quantification analysis (K) of apoptosis of mock or INHBA OE CT26 cells induced by IFN-γ (n = 3). The data are presented as the mean ±SEM. * p < 0.05; ** p < 0.01; *** p < 0.001; ns not significant by unpaired t test or one-way/two-way ANOVA followed by Tukey’s multiple comparisons test.

Subsequently, we tried to determine if INHBA represses IFN-γ response *in vitro*. Firstly, we assessed the phosphorylation level of STAT1 induced by IFN-γ. After IFN-γ stimulation for 1 or 2 hours, phosphorylation of STAT1 at Y701 significantly increased in mock cells, while at the same time points, INHBA overexpression inhibited IFN-γ induced STAT1 phosphorylation (Fig3E). Next, we investigated the expression of several IFN-γ-induced genes, namely *Cd274*, *Stat1*, *Cxcl9* and *Cxcl10*^31, 32, 33^. IFN-γ stimulation dramatically upregulated the expression of these genes in mock cells. And the upward trend was slowed down by *Inhba* overexpression in both CT26 and MC38 cells (Fig3F-3I, FigS6D-S6G). As a well-known tumor-killing factor, IFN-γ can also drive apoptosis of tumor cells^34^. Our *in vitro* apoptosis assay found that the sensitivity of tumor cells to IFN-γ-induced cytotoxicity was reduced by INHBA (Fig3J and 3K). In aggregate, IFN-γ signaling plays a crucial role in the crosstalk between INHBA and tumor immunity.

### INHBA suppresses IFNGR expression on tumor cells

The formation of IFN-γ receptor complex initiates the transmission of IFN-γ signaling^35^, so we first investigated if INHBA participates in regulating the IFN-γ receptor complex. The transcription levels of major components of IFN-γ receptor complex including IFNGR1, IFNGR2, JAK1 and JAK2 were detected and we found that INHBA negatively regulates the mRNA levels of all four genes both in cell lines and tumor tissues (Fig4A, 4B, 4E and FigS7A). By means of flow cytometry or western blotting in vitro, we noticed a drastic suppression of cellular IFNGR1 expression by tumor INHBA, while no alternation was observed in regard to expression of IFNGR2, JAK1 and JAK2 (Fig4C, 4D, FigS7B-S7C). Inverse correlation between tumor INHBA and IFNGR1 was also found in tumor tissues (Fig4F, 4G and FigS7D-S7E).

**Fig. 4.**
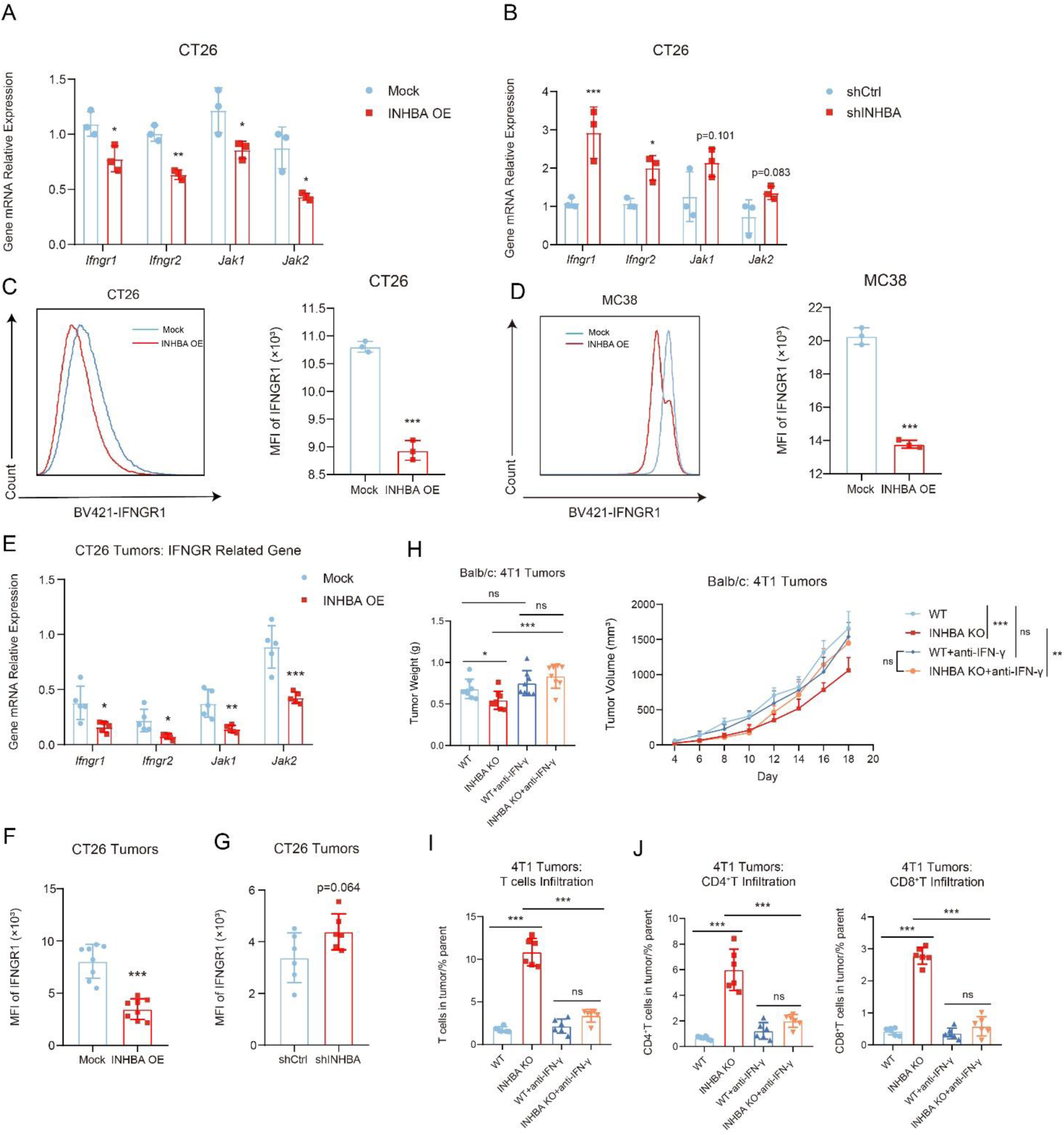
INHBA suppressed IFNGR expression on tumor. (**A** to **B**) Gene expression for *Ifngr1*, *Ifngr2*, *Jak1* and *Jak2* in mock or INHBA OE CT26 cells (A) and shCtrl or shINHBA CT26 cells (B; n = 3). (**C** to **D**) Representative histograms (left panel) and quantification analysis by flow cytometry (right panel) showing the effect of *Inhba* overexpression on IFNGR1 expression in CT26 (C) and MC38 cells (D; n = 3). (**E**) Gene expression for *Ifngr1*, *Ifngr2*, *Jak1* and *Jak2* in mock or INHBA OE CT26 tumors (n = 5). (**F** to **G**) Flow cytometric analysis showing the MFI of IFNGR1 in mock or INHBA OE CT26 tumors (F; n = 8) and shCtrl or shINHBA CT26 tumors (G; n = 6). (**H**) Tumor weights (left panel) and growth kinetics (right panel) of WT or INHBA KO 4T1 cells in Balb/c mice with or without treatment of anti-IFN-γ (n = 8). (**I**) Flow cytometric analysis showing the percentage of T cells infiltration in WT or INHBA KO 4T1 tumors with or without treatment of anti-IFN-γ (n = 6). (**J**) Flow cytometric analysis showing the percentage of CD4^+^ T and CD8^+^ T cells infiltration in WT or INHBA KO 4T1 tumors with or without treatment of anti-IFN-γ (n = 6). The data are presented as the mean ±SEM. * p < 0.05; ** p < 0.01; *** p < 0.001; ns not significant by unpaired t test or one-way ANOVA followed by Tukey’s multiple comparisons test.

However, the exact pathway by which INHBA affects IFNGR1 expression remains uncertain. Results of RNA-Seq showed that *Inhba* overexpression caused downregulation of multiple transcription factors of IFNGR1 (FigS7F). Further investigation is needed for the screening of core mediators. We then attempted to determine that tumor progression induced by INHBA depends on IFN-γ signaling in vivo. In *Inhba* knockout tumors, anti-IFN-γ treatment abolished tumor suppression and increased T cell infiltration (Fig4H-4I). To conclude, here, we suppose that the negative regulation of IFNGR1 by INHBA might account for its inhibition of the IFN-γ response.

### INHBA suppresses tumor PD-L1 expression via IFN-γ pathway

Clinically, tumor PD-L1 expression is an important marker for predicting the anti-PD-1/PD-L1 inhibitor effect; high PD-L1 expression endows patients with a higher objective response rate (ORR) ^7, 36^. As we described before, tumor derived INHBA presented a barrier to IFN-γ signaling transmission, for example, inhibiting the transcription level of IFN-γ induced PD-L1 (Fig3F). Taken together, we made the hypothesis that INHBA represses anti-PD-L1 efficacy by downregulating tumor PD-L1 expression.

Without IFN-γ stimulation, INHBA made no difference to the *Cd274* mRNA level (Fig5A and FigS8A). However, results of flow cytometry showed that *Inhba* overexpression hindered membrane PD-L1 expression induced by IFN-γ (Fig5B and FigS8B). Subsequently, in vivo models also implied that INHBA might negatively control tumor PD-L1 expression (Fig5C, Fig5D and FigS8C-S8D). Based on analysis of patients, PD-L1^+^ macrophages are more abundant in tumors than PD-L1^+^ cancer cells^37^. So we were also curious about whether Activin A could be secreted to impact PD-L1 expression on macrophages. Indeed, *Inhba* overexpression downregulated macrophage PD-L1 as well (FigS8E). Interestingly, we found that PD-L1 expression on tumor was elevated after atezolizumab treatment (Fig5E and 5F). It may be related to the activation of IFN-γ signaling, as *Inhba* overexpression abolished and *Inhba* knockdown further spurred PD-L1 elevation. The aforementioned data showed that T cells may be the major immune cell subset involved in INHBA regulation of oncogenesis. As IFN-γ is regarded as a weapon for T cells against cancer cells, we sought to confirm if INHBA inhibits the response to T cell-secreted IFN-γ. After co-culturing B16 with activated T cells for 24 h, we could see a stark increase in PD-L1 expression in B16 mock cells (Fig5G), but *Inhba* overexpression inhibited activated T cells induced PD-L1 elevation. What’s more, anti-IFN-γ treatment significantly repressed PD-L1 upregulation and eliminated the discrepancy between the mock and *Inhba* overexpression groups, implying that INHBA could diminish tumor PD-L1 expression induced by T cell derived IFN-γ.

**Fig. 5.**
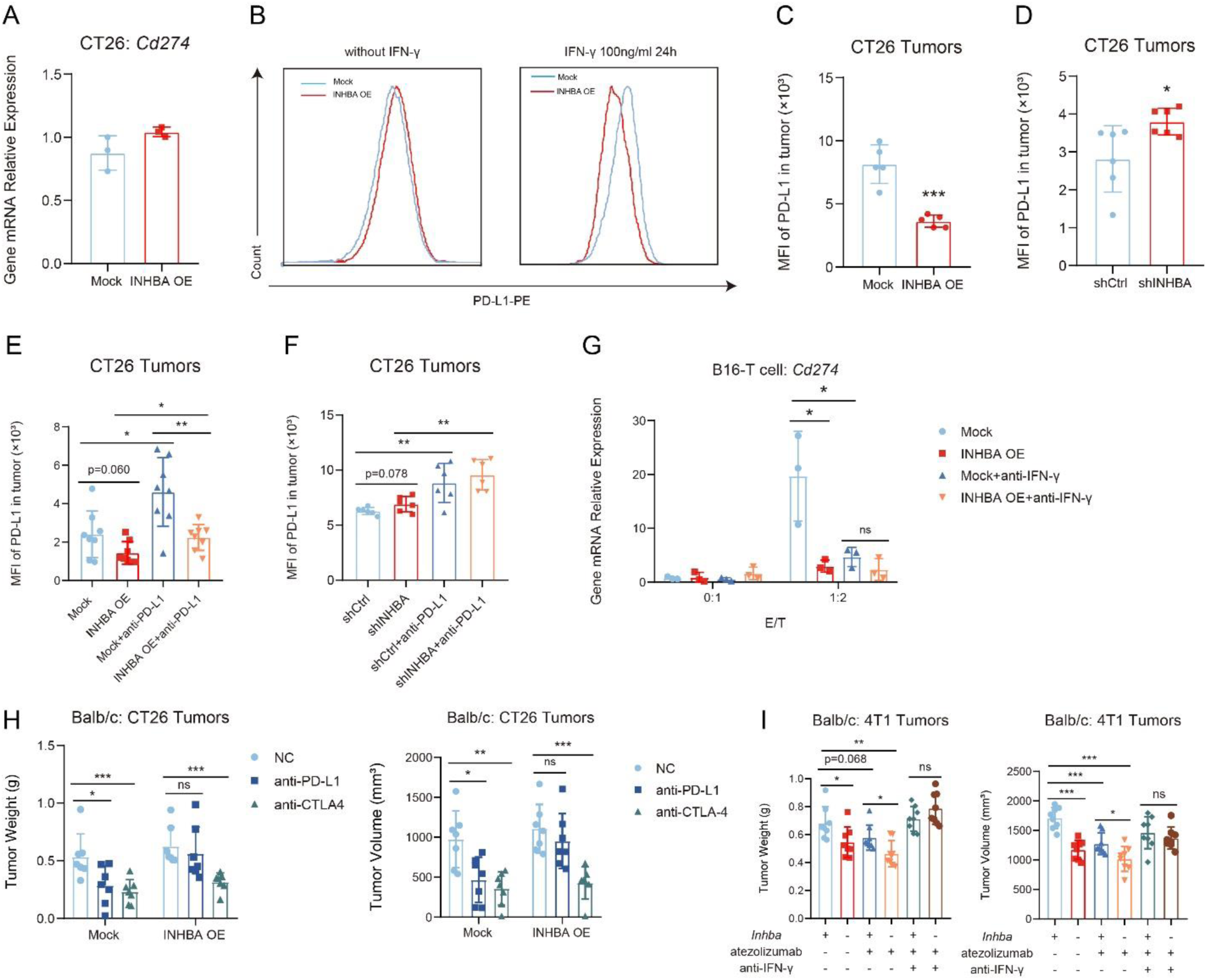
INHBA suppressed PD-L1 expression on tumor via IFN-γ pathway. (**A**) Gene expression for *Cd274* in mock or INHBA OE CT26 cells (n = 3). (**B**) Representative histograms of PD-L1 expression in mock or INHBA OE CT26 cells with treatment of mouse IFN-γ. (**C** to **D**) Flow cytometric analysis showing the MFI of PD-L1 in mock or INHBA OE CT26 tumors (C; n = 5) and shCtrl or shINHBA CT26 tumors (D; n = 6) (**E** to **F**) Flow cytometric analysis showing the MFI of PD-L1 in mock or INHBA OE CT26 tumors (E; n = 8) and shCtrl or shINHBA CT26 tumors (F; n = 6) with or without treatment of atezolizumab. (**G**) Gene expression of *Cd274* in mock or INHBA OE B16 cells co-cultured with activated splenic T cells with or without treatment of anti-IFN-γ (n = 3). (**H**) Tumor weights (left panel) and volumes (right panel) of mock or INHBA OE CT26 cells in Balb/c mice with non-treatment (NC), anti-PD-L1 or anti-CTLA4 treatment (n = 7). (**I**) Tumor weights (left panel) and volumes (right panel) of WT or INHBA KO 4T1 cells in Balb/c mice with non-treatment, atezolizumab or atezolizumab plus anti-IFN-γ (combination) (n = 8). The data are presented as the mean ±SEM and analyzed by unpaired t test or one-way/two-way ANOVA followed by Tukey’s multiple comparisons test.

Owing to the overall imbalance of T cell immunity caused by INHBA, we wonder if INHBA may result in resistance to other types of ICB therapy. As a result, mice bearing *Inhba* overexpression cells achieved a similar tumor repression rate to that bearing mock cells, indicating that INHBA induced resistance to ICB might be dependent on PD-L1 expression (Fig5H). To further validate IFN-γ signaling counts in INHBA-induced PD-L1 blockade resistance *in vivo*, we treated mice with 4T1 tumors, which poorly responses to atezolizumab, with IFN-γ antibody. As is expected, anti-IFN-γ abrogated the enhanced response to atezolizumab induced by *Inhba* knockout (Fig5I). These results suggest that by blocking T cell derived IFN-γ induced PD-L1 expression, INHBA induces resistance to anti-PD-L1 therapy.

### INHBA promotes tumor growth by suppressing secretion of CXCL9 and CXCL10

Next, we put efforts into revealing the concrete mechanism of how INHBA influences T cells in the tumor microenvironment. Based on our previous findings, INHBA can negatively regulate IFN-γ-induced transcription of *Cxcl9* and *Cxcl10* (Fig3G and 3H). CXCL9 and CXCL10, serving as IFN-γ induced chemokines, can lead to T cell recruitment to tumor tissues by binding to the CXCR3 receptor on T cells^38, 39^. After being treated with IFN-γ for 24 h, the supernatant of CT26 was collected and tested for CXCL9 and CXCL10 secretion by enzyme-linked immunosorbent assays (ELISAs). *Inhba* overexpression inhibited IFN-γ-induced secretion of CXCL10 (Fig6A). However, the concentration of CXCL9 did not reach the detection limit. Furthermore, we detected the release of CXCL9 and CXCL10 in tumor tissues. Similarly, *Inhba* overexpression significantly suppressed the secretion of both chemokines (Fig6B and 6C). It should be mentioned that CXCL10 had a substantially higher concentration than CXCL9, which could explain why the latter was undetectable in vitro. Additionally, tumor *Inhba* expression seemed to be positively correlated with CXCR3^+^ T cell infiltration *in vivo* (Fig6D-6G, FigS9A and S9C).

**Fig. 6.**
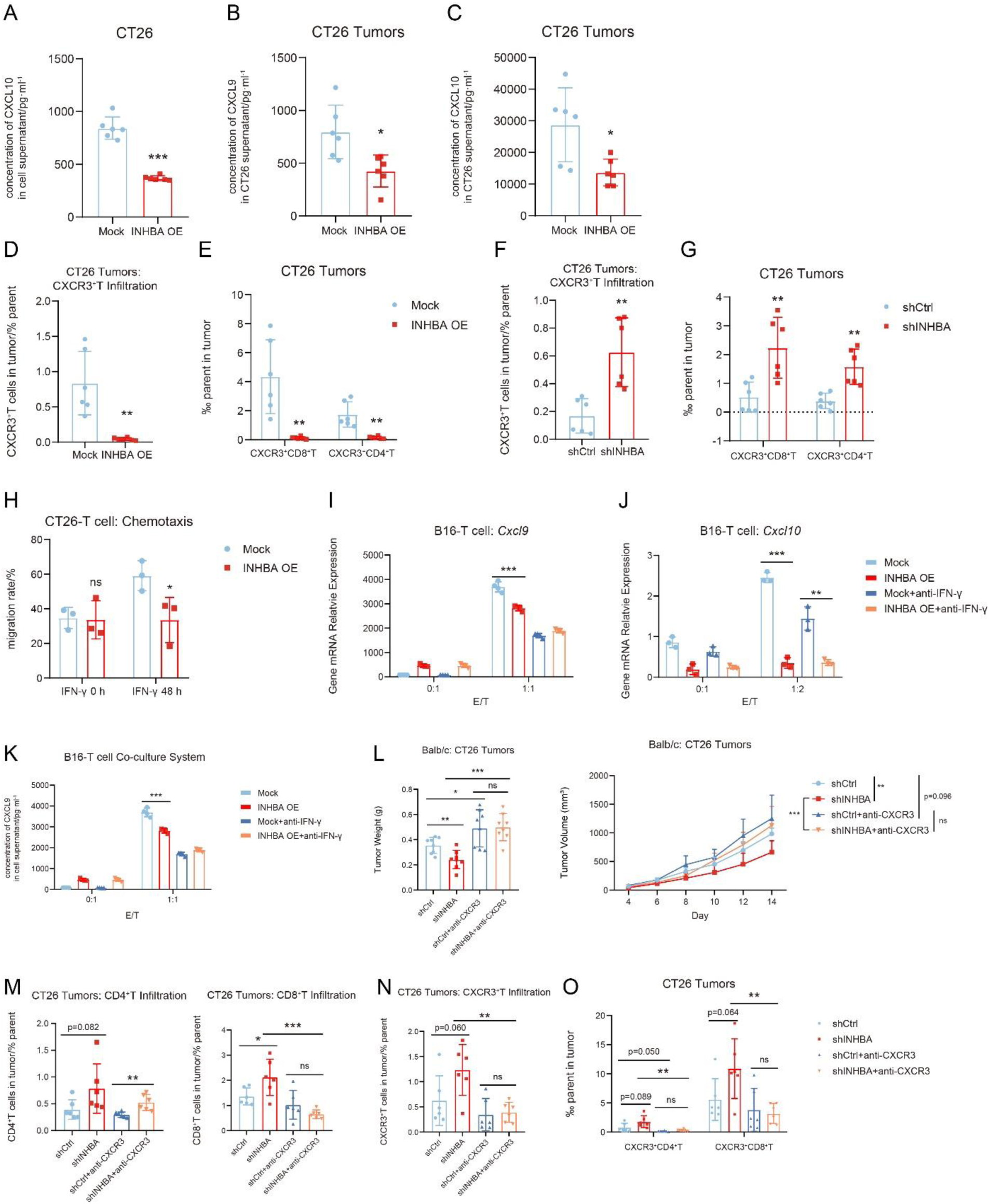
**INHBA promoted tumor growth by suppressing secretion of CXCL9 and CXCL10.** (**A**) The concentration of CXCL10 was determined in mock or INHBA OE CT26 cell supernatant after treatment of mouse IFN-γ (n = 6). (**B** to **C**) The concentration of CXCL9 (B) and CXCL10 (C) were determined in mock or INHBA OE CT26 tumor lysates (n = 6). (**D**) Flow cytometric analysis showing the percentage of CXCR3^+^ T cells infiltration in mock or INHBA OE CT26 tumors (n = 6). (**E**) Flow cytometric analysis showing the percentage of CXCR3^+^ CD4^+^ T and CXCR3^+^ CD8^+^ T cells infiltration in mock or INHBA OE CT26 tumors (n = 6). (**F**) Flow cytometric analysis showing the percentage of CXCR3^+^ T cells infiltration in shCtrl or shINHBA CT26 tumors (n = 6). (**G**) Flow cytometric analysis showing the percentage of CXCR3^+^CD4^+^T and CXCR3^+^CD8^+^T cells infiltration in shCtrl or shINHBA CT26 tumors (n=6). (**H**) Migration rate of activated T cells co-cultured with supernatant of mock or INHBA OE CT26 cells with or without treatment of mouse IFN-γ for 48 h (n = 3). (**I** to **J**) Gene expression of *Cxcl9* (I) and *Cxcl10* (J) in mock or INHBA OE B16 cells co-cultured with activated splenic T cells with or without anti-IFN-γ treatment (n = 3). (**K**) Concentration of CXCL9 in supernatant of mock or INHBA OE B16 cells co-cultured with activated splenic T cells with or without treatment of anti-IFN-γ (n = 3). (**L**) Tumor weights (left panel) and growth kinetics (right panel) of shCtrl or shINHBA CT26 cells in Balb/c mice with or without treatment of anti-CXCR3 (n = 8). (**M** to **O**) Flow cytometric analysis showing the percentage of CD4^+^ T (M; left), CD8^+^ T (M; right), CXCR3^+^ T (N), CXCR3^+^ CD4^+^ T and CXCR3^+^ CD8^+^ T cells (O) infiltration in shCtrl or shINHBA CT26 tumors with or without treatment of anti-CXCR3 (n = 6). The data are presented as the mean ±SEM. * p < 0.05; ** p < 0.01; *** p < 0.001; ns not significant by unpaired t test or one-way/two-way ANOVA followed by Tukey’s multiple comparisons test.

To ascertain if INHBA induced lower secretion of CXCL9 and CXCL10 directly results in reduced T cell chemotaxis, we co-cultured supernatants of cancer cells pre-treated with IFN-γ with activated T cells in a transwell system. Obviously, supernatants from *Inhba* overexpression cells obstructed T cell migration compared with that from mock cells (Fig6H). To support the speculation that attenuation in response to T cell-derived IFN-γ gives rise to decreased CXCL9 and CXCL10 secretion in *Inhba* overexpression cells, we directly contacted B16 with activated T cells. It is observable that INHBA hindered T cell induced transcriptional upregulation of *Cxcl9* and *Cxcl10*, along with the release of CXCL9 (Fig6I-6K). Notably, CXCL10 secretion was not detected in B16, which suggests that tumor heterogeneity exists in regard to the secretion of CXCL9 and CXCL10. IFN-γ blockade dampened the difference in chemokine secretion between the two groups, implying inhibition of T cell stimulated release of CXCL9 and CXCL10 by INHBA is at least partly dependent on IFN-γ. Moreover, treatment of IFN-γ antibody *in vivo* eliminated the effects of INHBA on CXCL9 and CXCL10 secretion and CXCR3^+^ T cell chemotaxis (FigS9D-S9F).

To consolidate that CXCL9/CXCL10/CXCR3 axis counts in INHBA-mediated tumor suppression, we treated CT26-bearing mice with an anti-CXCR3 antibody. Ablation of tumor inhibitory effect by *Inhba* knockdown was observed after anti-CXCR3 treatment, together with enhanced infiltration of CD8^+^ T and CXCR3^+^ T cells (Fig6L-6O). However, anti-CXCR3 scarcely affected total CD4^+^ T infiltration.

Above all, INHBA may promote tumor growth by inhibiting the secretion of CXCL9 and CXCL10 by tumor cells, thereby preventing T cell infiltration into the tumor site.

### Anti-activin A antibdy can suppress tumor growth and enhance responsiveness to anti-PD-L1 therapy

Given that the lack of INHBA in tumor drastically suppresses tumor progression and the responsive rate to anti-PD-L1 therapy, we suppos that specific therapeutic approaches targeting tumor INHBA/activin A might provide benefit to patients who are refractory to PD-L1 blockade. Currently, the major factor which retards the development of the activin A targeting strategy is a defect in specificity. A monoclonal antibody called garetosmab is regarded as the most specific Activin A inhibitor up to this point. It is now under a phase III clinical trial for the treatment of fibrodysplasia ossificans progressive (FOP). So far, rare cases have reported the application of garetosmab in cancer. Garetosmab shares similar binding affinity to Activin A receptor among human, mouse and monkey as Activin A is conserved across these species^40^.

Considering that our study mainly focused on the function of tumor endogenous INHBA, we adopted peritumoral injection for garetosmab (Fig7A). Garetosmab treatment showed a pronounced tumor inhibitory effect in CT26 model and facilitated the infiltration of total T cells, CD4^+^ T, CD8^+^ T and CXCR3^+^ T cells (Fig7B-7F). Subsequently, we combined garetosmab with atezolizumab to test if garetosmab could improve the efficacy of atezolizumab in the 4T1 tumor model, which is insensitive to anti-PD-L1 therapy. The results showed that the combinational treatment elicited a better tumor suppressive effect than garetosmab or atezolizumab alone (Fig7G). In addition, no matter whether combined with atezolizumab or not, garetosmab upregulated IFNGR1 and PD-L1 expression on tumor (Fig7H and 7I). Secretion of CXCL9 and CXCL10 was boosted by garetosmab or atezolizumab as well (Fig7J and 7K). Surprisingly, garetosmab also promoted CXCL9 secretion from macrophages in CT26 tumor, but without influence on that from DCs (FigS10A and S10B). All of the above results are consistent with our findings in the *Inhba* gene deficiency model, suggesting that garetosmab is a promising candidate to promote tumor regression and overcome resistance to anti-PD-L1.

**Fig. 7.**
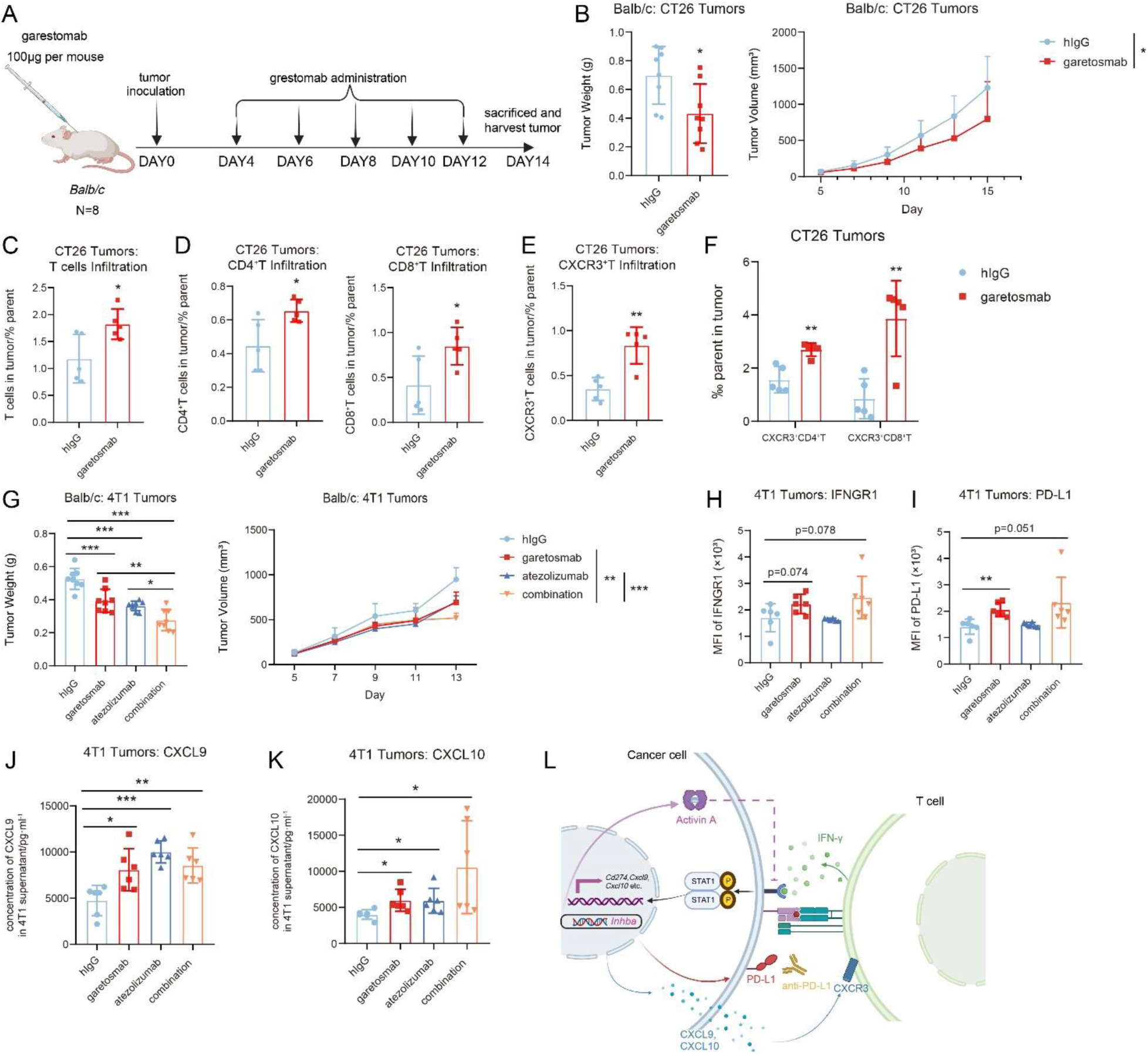
Anti-activin A antibody could suppress tumor growth and enhance responsiveness to anti-PD-L1 therapy. (**A**) Schematic of treatment schedules of garetosmab in mice bearing CT26 or 4T1 tumors. (**B**) Tumor weights (left panel) and growth kinetics (right panel) of CT26 cells in Balb/c mice treated with human IgG (hIgG) or garetosmab (n = 8). (**C** to **F**) Flow cytometric analysis showing the percentage of total T (C), CD4^+^ T (D left), CD8^+^ T (D right), CXCR3^+^ T (E), CXCR3^+^ CD4^+^ T and CXCR3^+^ CD8^+^ T cells (F) infiltration in CT26 tumors treated with hIgG or garetosmab (n = 5). (**G**) Tumor weights (left panel) and growth kinetics (right panel) of 4T1 cells in Balb/c mice treated with hIgG, garetosmab, atezolizumab or combination of garetosmab and atezolizumab (n = 8). (**H** to **I**) Flow cytometric analysis showing the MFI of IFNGR1 (H) and PD-L1 (I) in 4T1 tumors treated with hIgG, garetosmab, atezolizumab or combination (n = 6). (**J** to **K**) The concentration of CXCL9 (J) and CXCL10 (K) were determined in 4T1 tumor lysates treated with hIgG, garetosmab, atezolizumab or combination (n = 6). (**L**) The summary of role of tumor INHBA in regulating tumor immunity and anti-PD-L1 therapy. The data are presented as the mean ±SEM. * p < 0.05; ** p < 0.01; *** p < 0.001; ns not significant by unpaired t test or one-way/two-way ANOVA followed by Tukey’s multiple comparisons test.

## DISCUSSION

Our study provides potent evidence that indicates that tumor intrinsic INHBA can desensitize cancer cells to exogenous IFN-γ. On one hand, the blockade of T cell infiltration by reduced secretion of CXCL9 and CXCL10 promotes tumor progression; on the other hand, downregulated PD-L1 expression diminishes tumor responsiveness to PD-L1 antibodies (Fig7L). Application of an anti-activin A antibody also elicited considerable anti-tumor effects and potentiated the efficacy of PD-L1 blockade.

In the past decade, the advent of anti-PD-L1/PD-1 treatment has made a breakthrough in cancer immunotherapy. However, only a minority of patients acquired durable control of tumor progression^4^. Therefore, screening for key regulatory elements correlated to anti-PD-L1/PD-1 responsiveness will help overcome the failure of treatment. On the basis of clinical data, *Inhba* expression is elevated in tumors of melanoma patients unresponsive to anti-PD-L1/PD-1 axis^5, 30^.

IFN-γ plays a central role in cancer immune-editing. As a pleiotropic cytokine, IFN-γ impacts multiple aspects of tumor progression. Apart from directly inducing cancer cell apoptosis, IFN-γ also influences the functions of various tumor-related immune cells, such as promoting MHC molecule expression and tumor antigen presentation of antigen-presenting cells (APCs)^41^, directing macrophage polarization to M1 type^42^ and inducing differentiation of CD4^+^ T helper 1 cells (Th1) and CD8^+^ cytotoxic cells^43, 44^. In fact, during the tumor immune escape phase, IFN-γ elicits an opposite pro-tumor effect owing to the subsequent upregulation of PD-L1 ^45^. Based on the above effects, a predictable correlation exists between IFN-γ signaling imbalance and immune therapy resistance. Research has also corroborated that mutations in IFN-γ signaling core genes and the downregulation of IFNGR can lead to unresponsiveness to ICB ^46, 47^.

RNA-Seq of tumor tissues indicated that IFN-γ response signaling might play a key role in the relationship between INHBA and tumor immunity. Mechanically, INHBA may inhibit IFN-γ signaling by downregulating expression of IFNGR1 on tumor cells. As an IFN-γ induced product, tumor PD-L1 expression is regarded as an efficacy predictive biomarker for anti-PD-L1 treatment^48^. Moreover, the elevation of PD-L1 levels may be accompanied by a better therapeutic effect of PD-L1 antibody^49^. Our study found that INHBA might suppress tumor PD-L1 induced by IFN-γ, possibly from T cells, leading to the failure of PD-L1 blockade. The results of garetosmab treatment have deepened the idea that activin A is a potential therapeutic target for cancer immunotherapy. Our results re-emphasize the importance of IFN-γ signaling and tumor PD-L1 expression in anti-PD-L1 therapy.

Intratumoral immune cell infiltration level is also a potential anti-PD-L1/PD-1 efficacy influencing factor. According to TCGA datasets, upregulation of *Inhba* can be observed in diverse types of cancers and is broadly linked to poor prognosis, along with reduced infiltration of CD4^+^ T and CD8^+^ T cells. Few studies have shed light on how tumor intrinsic activin A impacts tumor immunity. It was reported that overexpression of *Inhba* in melanoma triggered impaired infiltration and cytotoxicity of CD8^+^T cells^27, 50^. Despite that, the specific regulatory mechanism is unclear, and it is insufficient to uncover the general role of activin A in tumor immunity as the research context is confined to melanoma.

Notably, deficient infiltration of CD4^+^ T and CD8^+^ T cells induced by INHBA was observed among all of our subcutaneous tumor models. IFN-γ can also induce secretion of chemokines, including CXCL9, CXCL10 and CXCL11^51^. Our research focused on CXCL9 and CXCL10 due to the controversial impact of CXCL11 on immune modulation^52, 53^. Myeloid cells and tumor cells can both release CXCL9 and CXCL10 in response to IFN-γ to recruit CXCR3^+^ T cells^54^.

Pinjusic, K et al. showed that in melanoma, activin A could inhibit CXCL9 and CXCL10 secretion from myeloid cells^27^. Instead, our findings established that INHBA could also decrease tumor-derived CXCL9 and CXCL10 via impairing IFN-γ responses. Anti-tumor function of INHBA was ablated after CXCR3 blockade, which further supported that INHBA depends on CXCL9 and CXCL10 to influence tumor growth. These results hint at the INHBA-IFNGR1-CXCL9/CXCL10 axis as a potential mechanism for INHBA-mediated inhibition of intratumoral infiltration of immune cells, which in turn further exacerbates anti-PD-L1 treatment resistance.

For sure, our present study has some limitations. First of all, we mainly discussed how tumor cells are affected by tumor intrinsic INHBA in response to IFN-γ. However, as a secreted cytokine, it is not yet clear whether tumor activin A exerts a similar effect on other cells in TME via paracrine, for example, inhibiting IFN-γ induced MHC expression or chemokine secretion in APCs. Besides, increasing evidence has demonstrated that myeloid PD-L1 levels seem to be associated with anti-PD-L1 efficacy as well^55^. We also preliminarily found that overexpression of INHBA may simultaneously inhibit PD-L1 expression and CXCL9 secretion in macrophages, but systematic validation is required. Secondly, with regard to the regulation of IFNGR1 expression, we did not explored it in depth, but indeed, we found that multiple transcription factors were downregulated in the context of INHBA overexpression. Additional efforts are needed to elucidate the concrete pathway. At last, to highlight the function of tumor derived INHBA, we administrated garetosmab in a peritumoral way. We will develop agents with better tumor-targeting ability, such as activin A/PD-L1-bispecific antibody, to reach optimal therapeutic effects.

In summary, our work validated the tumor intrinsic INHBA as a tumor immunity regulator that affects IFNGR1 expression to interfere with the IFN-γ-PD-L1 axis and the IFN-γ-CXCL9/CXCL10 axis to compromise immune cell infiltration and anti-PD-L1 therapy efficacy. The potential of INHBA as a target for tumor immunotherapy and in combination with anti-PD-L1/PD-1 antibodies has been strongly verified.

## METHODS

### Mice

Female (Six-to eight-week-old) BALB/c and C57BL/6 mice were purchased from the Shanghai Slack. Balb/c nu/nu mice were purchased from Vital River Laboratory. All mice were maintained under specific pathogen-free (SPF) conditions in the animal facility of the Shanghai Institute of Materia Medica, Chinese Academy of Sciences (SIMM). Animal care and experiments were performed in accordance with protocols approved by the Institutional Laboratory Animal Care and Use Committee (IACUC).

### Cell lines

The mouse colon carcinoma line CT26, MC38, melanoma cell line B16, breast cancer cell line 4T1, and HEK-293T were all obtained from 2019 to 2023 and passed short tandem repeat (STR) analysis and mycoplasma testing before use. B16 and HEK293T cells were cultured in Dulbecco’s Modified Eagle Medium (DMEM; MA0212, Meilunbio) supplemented with 1% penicillin/streptomycin (P/S; 15140122, Gibco) and 10% heat-inactivated fetal bovine serum (FBS; 10100147, Life Technologies). 4T1, MC38 and CT26 cells were cultured in RPMI1640 medium (MA0215; Meilunbio) containing 1% P/S and 10% FBS. The number of cell passages was controlled within 15 passages. Other stable cell lines were constructed by the lentivirus system and cultured under the same conditions as the parental cells (see *Construction of stable cell lines*). All cells were cultured at 37°C in a 5% CO2 humidified atmosphere.

### Plasmids and viruses

pSPAX2 (VT1444; Youbio) and pMD2.G (VT1443; Youbio) were used to package lentiviruses. pLVX-puro (VT1465; Youbio) was used to package vacant vector and pLVX-mINHBA-puro (G132770; Youbio) was used for INHBA overexpression. pLV-EGFP-Scramble_shRNA (VB010000-0009mxc; VectorBuilder) was used to express package vacant vector and pLV-EGFP-shInhba (VB900048-2616uaf; VectorBuilder) was used for INHBA knockdown. shRNA target sequences: mouse INHBA, 5′-GGCCGAGGAAATGGGCTTAAA-3′.

### Tumor models using transgenic tumor cells

For all the subcutaneous models, mice were randomly grouped and injected subcutaneously with 5×10^5^-1×10^6^ tumor cells in the right forelimb. Tumor volume was measured and recorded every two days. Tumor volume was calculated as tumor volume (mm^3^) = tumor length ×width ×width/2, and tumor growth curves were plotted. At least four time points after tumor inoculation, the mice were sacrificed and the tumors harvested for weighing, photographing, or other purposes. For survival studies, euthanasia endpoints were set as tumors larger than 1500 mm^3^ or larger than 20 mm in length or mice with significant weight loss.

### Construction of stable cell lines

Various INHBA knockdown or overexpression cell lines, were constructed using the lentiviral system. The corresponding shRNA is listed above (see *Plasmids and viruses*). Briefly, the gene of interest was subcloned into lentiviral expression vector (pLVX-puro or pLV-EGFP-Scramble_shRNA; 7 μg), which was then transfected using Lipofectamine 3000 (L3000150; Thermo Fisher) into HEK293T cells (10 cm cell culture dish; 60% cell confluency) with the package plasmids (pSPAX2 and pMD2.G; 4.2 μg/each). Three days later, the virosomes were collected by lentivirus concentration kit (GM-040801; Genomeditech) and inflected into target tumor cells. The stable cells were obtained after antibiotic puromycin (MA0318; Meilunbio) resistance selection and were identified by RT-qPCR, ELSIA, Western Blot or flow cytometry.

### RNA isolation and RT-qPCR analysis

Total RNA of cells or tumor tissues was isolated with TRIzol (10606ES60; Yeasen) and cDNA was generated with an RT reagent kit (RK20428; ABclonal). Quantitative PCR was carried out with 2× SYBR Green Fast qPCR Mix (RK21203; ABclonal) to assess target mRNA expression. The primers were obtained from GENEWIZ, China. The sequences of the primers used in this study were listed in **TableS1**. All reactions were performed at least in triplicate to determine the average Ct for each gene. GAPDH was chosen as an internal control, and the relative expression of each gene was calculated according to the 2^−ΔΔCt^ method.

### TCGA analysis

Clinical pan-cancer *INHBA* mRNA expression data were obtained from the The Cancer Genome Atlas (TCGA) datasets (https://portal.gdc.cancer.gov/) by GEPIA software. The number of tumor or normal samples is indicated in the figures. Expression differences, prognostic correlation, and immunological correlation were analyzed by GEPIA software.

### In vivo treatment studies

Studies of anti-CTLA-4 (BE0032, BioXcell) therapy in mice bearing CT26 and B16 tumors, atezolizumab (CHA083, Sanyou Bio) therapy in mice bearing CT26, 4T1, MC38 and B16 tumors, garetosmab (CHA401; Sanyou Bio), human IgG4 (COA001; Sanyou Bio), anti-CXCR3 (BE0249, BioXcell) and anti-IFN-γ (BE0055, BioXcell) therapy in mice bearing CT26 and 4T1 tumors are consistent with the descriptions above. The administration methods of each drug are shown in the respective flow charts in the figures.

### Immune cell depletion study

Mice were randomly divided and injected with tumor cells subcutaneously, which was defined as day 0. CD4^+^ T cells were deleted by intraperitoneal (i.p.) injection of 150 μg/each anti-CD4 (BE0119, BioXcell) on day -2, day 0, day 4 and day 8. CD8^+^ T cells were deleted by i.p. injection of 150 μg/each anti-CD8α (BE0061, BioXcell) on day -2, day 0, day 4 and day 8. The deletion effect was identified by flow cytometry (see *Immunophenotyping analysis*).

### Immunophenotype analysis

For immunotyping of intratumoral cells, tumor tissues were digested by collagenase IV (40510ES60, Yeasen) and hyaluronidase (20426ES60, Yeasen) and were filtered through 75 micron nylon mesh (7061011, Dakewe) into single cell suspensions and then were subjected to cell extraction using lymphocyte separation medium (7211011, Dakewe) for sorting tumor-infiltrating lymphocytes (TILs) or were subjected to erythrocyte removal by red blood cell lysis buffer (40401ES60, Yeasen) for non-lymphocyte staining. Then the cells were blocked with 4% FBS and anti-CD16/CD32 (553141, BD Biosciences), incubated with surface marker antibodies for 20 minutes at 4℃ and then permeabilized with BD Cytofix/Cytoperm buffer (554714) before intracellular labeling antibodies were added for 30 minutes at 4℃. For staining of cytokines, 1×10^6^ cells were stimulated with 2 μL leukocyte activation cocktail (550583, BD Biosciences) in 1 mL complete medium for 4 hours at 37°C in 5% CO2. Flow cytometry analysis was performed using ACEA NovoCyte and data processing was done through NovoExpress software (version 1.6.1). Antibody staining was performed following the manufacturer’s recommendations. Please refer to **TableS2** for information about the antibodies used. For immunotyping of splenocytes, single-cell suspensions were obtained by grinding tissues and were subject to erythrocyte removal. The other steps are the same as above.

### Western blot analysis

For the analysis of INHBA affecting activation of JAK-STAT1 signaling pathway in tumor cells, 5×10^5^ cells per well (6 well plate) were stimulated with 100 ng/mL of IFN-γ (Z02916, Genscript) for 0 h, 1 h and 2 h. Afterwards, cells were lysed by RIPA lysis buffer (P0013C, Beyotime) with PMSF (ST507, Beyotime) (the ratio of the two reagents is 100:1) on the ice. Proteins were separated by SDS-PAGE and then transferred to 0.45 μm PVDF membrane (IPVH00010, Merck Millipore). To keep the amounts of protein equally, β-tubulin was used as a loading control.

### Cell proliferation assay

For the cell proliferation assay, 5×10^3^ tumor cells were seeded in 96-well plates. CCK8 (40203ES60, Yeasen) was added at 0 h, 24 h, 48 h, 72 h, respectively. And the signal was read at OD 450 nm by an automatic microplate reader SpectraMax (Molecular Devices).

### Apoptosis assay

For the apoptosis assay, 2×10^5^ tumor cells were pre-treated with 200 ng/mL of IFN-γ for 0h and 12h in 12 well plate. Subsequently, the cells were harvested and washed twice with pre-cooled PBS, stained with annexin V-FITC and PI using an Apoptosis Detection Kit (40302ES20, Yeasen) at 25℃ for 15 min. The percentage of apoptosis cells was determined by flow cytometry.

### Isolation and activation of mouse spleen-derived CD3^+^ T cells

After isolation of mouse splenocytes as descripted above (see *Immunophenotyping analysis*), CD3^+^ T cells were isolated by magnetic bead sorting (Mouse T Cell Isolation Kit, 19851, Stemcell). For the activation of T cells, 2×10^6^ cells/well (6-well plate) were cultured with 1μg/mL of anti-CD3 mAb (553057, BD Biosciences) plus 10 ng/mL of mouse IL-2 (Z02764, Genscript) in 2 mL complete RPMI1640 medium for 72 h.

### In vitro T cell migration assays

For the analysis of the effect of INHBA on T cell migration driven by IFN-γ-induced secretion of CXCL9 and CXCL10, 5×10^5^ mock or INHBA OE tumor cells were treated with or without 100 ng/mL IFN-γ for 48 h. For the T cell migration assay, 1 mL conditioned media from tumor cells descripted above was added into the bottom chamber of a transwell device with a PET membrane with a 3 μm pore size (14322, LABSELECT), and the top chamber contained 1×10^5^ activated CD3^+^ T cells in 100 μL complete RPMI1640 medium.

After the co-culture for 24 h, the media of both the upper and the bottom chamber was collected and centrifuged at 13,000x g for 20 min at 4°C respectively. Afterwards, the supernatant was discarded and PBS was used to resuspend cells. Cell numbers were counted using Cell counting analyzer (BodBoge) and the migration rate was calculated as migration rate (%) = [number of cells in the bottom chamber/ (number of cells in the bottom chamber + number of cells in the upper chamber)] ×100%.

### ELISAs

For analysis of INHBA, CXCL9 and CXCL10 production by tumor cell lines, 5×10^5^ mock or INHBA OE cells per well were pre-treated with 100 ng/mL of IFN-γ for 24 h in a 6-well plate. The supernatant was collected and the concentration was quantified with ELISA kits as follows: INHBA (JL34309, Jianglaibio), CXCL9 (JL20226, Jianglaibio) and CXCL10 (JL13372, Jianglaibio). For analysis of CXCL9 and CXCL10 production in tumor tissues, 20 mg tumor tissues were lysed with PBS supplemented with PMSF. After centrifugation at 10,000x g for 10 min at 4 ℃, the supernatant was collected and the concentration of these cytokines was quantified with ELISA kits as follows: CXCL9 (JL20226, Jianglaibio) and CXCL10 (JL13372, Jianglaibio).

### Co-culture assay

For the T cells–B16 co-culture study, 2×10^5^ of B16 cells were seeded in 12-well plates 8 h in advance. T cells were isolated and activated (see *Isolation and activation of mouse spleen-derived CD3+ T cells*), then added to the wells at a ratio of 0:1, 1:2 and 1:1, respectively. After 24 h, T cells were removed by PBS washing and the supernatant was used for cytokine detection by ELISAs using the following kits for CXCL9 (JL20226, Jianglaibio) and CXCL10 (JL13372, Jianglaibio). mRNA expression of PD-L1, CXCL9 and CXCL10 of B16 after co-cultured with T cells was quantified by RT-qPCR (see *RNA isolation and RT-qPCR analysis*).

### Adoptive transfusion of immune cells

For adoptive transfusion of TILs, donor mice were inoculated with CT26 (5×10^5^) or 4T1 (1×10^6^) tumor cells seven days before the recipient Balb/c nu/nu mice. On day 7 of tumor growth in recipient mice, the donor mice were sacrificed and tumor tissue was obtained. Under aseptic conditions, TILs were isolated. TILs (5×10^6^) were peritumorally injected into recipient mice. Tumor growth kinetics, volumes and weights were measured.

### Other In vitro cell assays

For the analysis of the effect of INHBA on PD-L1 expression in tumor cells, mock or INHBA overexpression CT26 cells were directly analyzed for PD-L1 expression (BV650 Rat Anti-Mouse CD274; 740614, BD Bioscience) by flow cytometry, or after being induced by 100 ng/mL of IFN-γ for 24 h.

For the analysis of the effect of INHBA on IFNGR1 or IFNGR2 expression in tumor cells, mock or INHBA overexpression tumor cells (CT26 and MC38) and shCtrl or shINHBA CT26 cells were analyzed for IFNGR1 (BV421 Rat Anti-Mouse CD119; 740032, BD Bioscience) and IFNGR2 expression (Hamster Anti-Mouse IFNGR2 Monoclonal antibody; MAB773, R&D systems) by flow cytometry. For IFNGR2 detection, AF488 anti-Hamster IgG (A78963, Thermo Fisher) was used as the second antibody for IFNGR2 detection.

### RNA-Seq

About 30 mg of fresh tumor tissues were obtained from sacrificed CT26 tumor-bearing mice. RNA isolation, transcriptome libraries construction, sequencing and basic data analysis were conducted by BGI. Based on the RNA-seq raw data, differential expression was evaluated with DESeq. A fold-change of 2:1 or greater and a false discovery rate (FDR)-corrected p-value of 0.05 or less were set as the threshold for differential genes. Immune signature scores are defined as the mean log_2_(fold-change) among all genes in each gene signature list from Gene Set Enrichment Analysis (GSEA) datasets. Cell infiltration within tumor tissues was estimated by xCell (https://xcell.ucsf.edu/).

### Statistical analyses

The *in vivo* experiments were randomized but the researchers were not blinded to allocation during experiments and outcome analysis. Statistical analysis was performed using GraphPad Prism 8 Software. A Student’s t test was used for comparison between the two groups. Multiple comparisons were performed using one-way/two way ANOVA followed by Tukey’s multiple comparisons test. Survival analysis was performed by the Kaplan–Meier method. Detailed statistical methods and sample sizes in the experiments are described in each figure legend. No statistical methods were used to predetermine the sample size. All statistical tests were two-sided and P-values < 0.05 were considered to be significant. ns not significant; *p < 0.05; **p < 0.01; ***p < 0.001.

## Data availability

The raw sequence data reported in this paper could be found in **Supplementary File 2** and are publicly available as of the date of publication. This paper analyzes existing, publicly available data. These accession numbers for the datasets are listed in **TableS2**. Correspondence and requests for materials should be addressed to L.Gong. or Y.L.

## Acknowledgments

All the figures were created with BioRender.com. This work was supported by Foundation of Shanghai Science and Technology Committee (No.22S11902100), Zhongshan Municipal Bureau of Science and Technology (No. 2020SYF08), the Department of Science and Technology of Guangdong Province (No. 2019B090904008 and No. 2021B0909050003), and the Strategic Priority Research Program of the Chinese Academy of Sciences (No. XDA 12050305).

## Author contributions

Conceptualization, L.Gong, Y.L. and F.L.; Methodology, Y.L., Y.T., X.Y. and F.L.; Formal Analysis, F.L.; Investigation, F.L., L.Gu., R.C., S.C., J.L. and Y.S.; Resources, N.L. and Y.L.; Writing – Original Draft, F.L.; Writing – Review & Editing, L.Gong and Y.L.; Supervision, L.Gong; Funding Acquisition, L.Gong.

