## Supplementary file 1 for "INHBA/Activin A promotes tumor growth and induces resistance to anti-PD-L1 therapy by suppressing IFN-γ signaling"

**This file contains:**

TableS1. Primer sequences for RT-PCR analysis.

TableS2. List of key resources.

FigS1. Effect of INHBA on tumor growth.

FigS2. Correlation between tumor INHBA and immune cell infiltration.

FigS3. Effect of INHBA on tumor immunity.

FigS4. Validation of immune cell depletion efficacy.

FigS5. Effect of INHBA on anti-PD-L1 resistance.

FigS6. Effect of INHBA on IFN-γ response.

FigS7. Effect of INHBA on expression of IFN-γ receptor complex.

FigS8. Effect of INHBA on tumor PD-L1 expression.

FigS9. Effect of INHBA on CXCL9/CXCL10/CXCR3 axis.

FigS10. Effect of garetosmab on CXCL9 secretion from myeloid cells.

**TableS1. Primer sequences for RT-PCR analysis.**

| **Gene** | **Primer** | **Sequence (5'→3')** |
| --- | --- | --- |
| *Gapdh* | FW | TCTCCTGCGACTTCAACA |
|  | RV | TGGTCCAGGGTTTCTTACT |
| *Inhba* | FW | GGGGAGAACGGGTATGTGGA |
|  | RV | CCTGACTCGGCAAAGGTGAT |
| *Jak1* | FW | CTCTCTGTCACAACCTCTTCGC |
|  | RV | TTGGTAAAGTAGAACCTCATGCG |
| *Jak2* | FW | TTGTGGTATTACGCCTGTGTATC |
|  | RV | ATGCCTGGTTGACTCGTCTAT |
| *Ifngr1* | FW | TACAGGTAAAGGTGTATTCGGGT |
|  | RV | ACCGTGCATAGTCAGATTCTTTT |
| *Ifngr2* | FW | TCCTCGCCAGACTCGTTTTC |
|  | RV | GTCTTGGGTCATTGCTGGAAG |
| *Cd274* | FW | GCTCCAAAGGACTTGTACGTG |
|  | RV | TGATCTGAAGGGCAGCATTTC |
| *Cxcl9* | FW | GGAGTTCGAGGAACCCTAGTG |
|  | RV | GGGATTTGTAGTGGATCGTGC |
| *Cxcl10* | FW | CCAAGTGCTGCCGTCATTTTC |
|  | RV | GGCTCGCAGGGATGATTTCAA |
| *Stat1* | FW  RV | TCACAGTGGTTCGAGCTTCAG  GCAAACGAGACATCATAGGCA |

**TableS2. List of key resources.**

| REAGENT or RESOURCE | SOURCE | IDENTIFIER |
| --- | --- | --- |
| Antibodies | | |
| Anti-mouse INHBA (clone ARC1177) | ABclonal | Cat# A5232; RRID:AB_2863495 |
| Anti-mouse STAT1 (clone ARC0042) | ABclonal | Cat#A19563; RRID:AB_2862669 |
| Anti-mouse Phospho-STAT1-Y701 (clone ARC0049) | ABclonal | Cat#AP0054; RRID:AB_2863803 |
| Anti-mouse GAPDH (clone ARC50888) | ABclonal | Cat#A19056; RRID:AB_2862549 |
| Anti-mouse β-Tubulin (clone ARC0203) | ABclonal | Cat#A12289; RRID:AB_2861647 |
| Anti-mouse JAK1 (clone ARC0434) | ABclonal | Cat# A11963 |
| Anti-mouse JAK2 (clone ARC0108) | ABclonal | Cat# A19629;  RRID:AB_2862706 |
| APC-Cy7 anti-mouse CD45 (clone 30-F11) | BD Biosciences | Cat#561037; RRID:AB_10563075 |
| APC-Cy7 anti-mouse CD3e (clone 145-2C11) | BD Biosciences | Cat#561042; RRID:AB_2034003 |
| BV421 anti-mouse CD4 (clone RM4-4) | BD Biosciences | Cat#740008; RRID:AB_2739780 |
| BV510 anti-mouse CD8a (clone 53-6.7) | BD Biosciences | Cat#563068; RRID:AB_2687548 |
| BV650 anti-mouse CD274 (clone MIH5) | BD Biosciences | Cat#740614; RRID:AB_2740313 |
| PE anti-mouse CD274 (clone MIH5) | BD Biosciences | Cat# 558091; RRID:AB_397018 |
| BV421 anti-mouse CD11b (clone M1/70) | BD Biosciences | Cat#562065; RRID:AB_11152949 |
| APC anti-mouse CD183 (clone CXCR3-173) | BD Biosciences | Cat#562266; RRID:AB_11153500 |
| BV421 anti-mouse CD119 (clone GR20) | BD Biosciences | Cat#740032; RRID:AB_2739804 |
| BV786 anti-mouse IFN-γ (clone XMG1.2) | BD Biosciences | Cat#563773; RRID:AB_2738419 |
| PE-CF594 anti-mouse IL-4 (clone 11B11) | BD Biosciences | Cat#562450; RRID:AB_2737616 |
| PE anti-mouse CD278 (clone 7E.17G9) | BD Biosciences | Cat# 552146; RRID:AB_394349 |
| BV650 anti-mouse IL-10 (JES5-16E3) | BD Biosciences | Cat#564083; RRID:AB_2738583 |
| APC anti-mouse CD86 (clone GL1) | BD Biosciences | Cat#561964; RRID:AB_10898000 |
| Anti-mouse CD3e (clone 145-2C11) | BD Biosciences | Cat# 553057; RRID:AB_394590 |
| Anti-mouse CD16/CD32 (clone 2.4G2) | BD Biosciences | Cat# 553141; RRID:AB_394656 |
| PE anti-mouse CXCL9 (clone MIG-2F5.5) | eBioscience | Cat# 12-3009-80  RRID: AB_891582 |
| AF488 anti-Hamster IgG | ThermoFisher Scientific | Cat#A78963; RRID:AB_2925786 |
| AF647 anti-Rabbit IgG | Abcam | Cat#ab150083; RRID:AB_2714032 |
| FITC anti-mouse CD206 (clone C068C2) | Biolegend | Cat#141704; RRID:AB_10901166 |
| BV605 anti-mouse CD152 (clone UC10-4B9) | Biolegend | Cat#106323; RRID:AB_2566467 |
| PE-Cy7 anti-human/mouse Granzyme B (clone QA16A02) | Biolegend | Cat#372214; RRID:AB_2728381 |
| PE-Cy7 anti-mouse F4/80 (clone BM8) | Biolegend | Cat# 123114; RRID:AB_893478 |
| Anti-mouse IFNGR2 (clone mob47) | R&D systems | Cat#MAB773; RRID:AB_2248682 |
| InVivoMAb anti-mouse CD4 (clone YTS 191) | BioXcell | Cat#BE0119; RRID:AB_10950382 |
| InVivoMAb anti-mouse CD8a (clone 2.43) | BioXcell | Cat#BE0061; RRID:AB_1125541 |
| InVivoMAb anti-mouse IFN-γ (clone XMG1.2) | BioXcell | Cat#BE0055; RRID:AB_1107694 |
| InVivoMAb anti-mouse CTLA-4 (clone UC10-4F10-11) | BioXcell | Cat#BE0032; RRID:AB_1107598 |
| InVivoMAb anti-mouse CXCR3 (clone CXCR3-173) | BioXcell | Cat#BE0249; RRID:AB_2687730 |
| Anti-INHBA / Activin A reference antibody garetosmab | Sanyou Bio | Cat# CHA401 |
| Anti-B7-H1 / PD-L1 / CD274 reference antibody atezolizumab | Sanyou Bio | Cat# CHA083 |
| Anti-human IgG4 reference antibody | Sanyou Bio | Cat# COA001 |
| Bacterial and virus strains | | |
| Stbl3 | Absin | Cat# abs60045 |
| Chemicals, peptides, and recombinant proteins | | |
| Mouse IFN-γ | Genscript | Cat# Z02916 |
| Mouse IL-2 | Genscript | Cat# Z02764 |
| Leukocyte Activation Cocktail | BD Biosciences | Cat#550583;  RRID: AB_2868893 |
| Lipofectamine 3000 | ThermoFisher Scientific | Cat# L3000150 |
| Puromycin | Meilunbio | Cat# MA0318 |
| TRIzol | Yeasen | Cat# 10606ES60 |
| Collagenase IV | Yeasen | Cat# 40510ES60 |
| Hyaluronidase | Yeasen | Cat# 20426ES60 |
| Red blood cell lysis buffer | Yeasen | Cat# 40401ES60 |
| Lymphocyte separation medium | Dakewe | Cat# 7211011 |
| RIPA lysis buffer | Beyotime | Cat# P0013C |
| PMSF | Beyotime | Cat# ST507 |
| 2X SYBR Green Fast qPCR Mix | ABclonal | Cat# RK21203 |
| Critical commercial assays | | |
| Cytofix/Cytoperm Soln Kit | BD Biosciences | Cat#554714; RRID:AB_2869008 |
| CCK8 Kit | Yeasen | Cat# 40203ES60 |
| Annexin V-FITC/PI Apoptosis Detection Kit | Yeasen | Cat# 40302ES20 |
| Mouse T Cell Isolation Kit | Stemcell | Cat# 19851 |
| Mouse CXCL9 ELISA Kit | Jianglaibio | Cat# JL20226 |
| Mouse CXCL10 ELISA Kit | Jianglaibio | Cat# JL13372 |
| Mouse INHBA ELISA Kit | Jianglaibio | Cat# JL34309 |
| Lentivirus Concentration Kit | Genomeditech | Cat# GM-040801 |
| RT Reagent Kit | ABclonal | Cat# RK20428 |
| Deposited data | | |
| RNA-Seq data of melanoma | Hugo et al. | GEO: GSE78220 |
|  | Riaz et al. | GEO: GSE91061 |
| TCGA pan-cancer data | The Cancer Genome Atlas (TCGA) | https://portal.gdc.cancer.gov/ |
| RNA-Seq data | This paper |  |
| Experimental models: Cell lines | | |
| CT26 | Procell | Cat# CL-0071 |
| B16 | Procell | Cat# CL-0029 |
| 4T1 | Procell | Cat# CL-0007 |
| HEK-293T | Procell | Cat# CL-0005 |
| MC38 | Yuchun Biology | Cat# CM0042 |
| Experimental models: Organisms/strains | | |
| C57/BL6J mice | Shanghai Slack | N/A |
| BALB/c mice | Shanghai Slack | N/A |
| BALB/c nu/nu mice | Vital River Laboratory | N/A |
| Oligonucleotides | | |
| Primers used for qRT-PCR (see Table S1) | This paper | N/A |
| Recombinant DNA | | |
| pSPAX2 | Youbio | Cat# VT1444 |
| pMD2.G | Youbio | Cat# VT1443 |
| pLVX-puro | Youbio | Cat# VT1465 |
| pLVX-mINHBA-puro | Youbio | Cat# G132770 |
| pLV[shRNA]-EGFP/Puro-U6>Scramble_shRNA | VectorBuilder | Cat# VB010000-0009mxc |
| pLV[shRNA]-EGFP:T2A:Puro-U6>mInhba | VectorBuilder | Cat# VB900048-2616uaf |
| Software and algorithms | | |
| ImageJ | Schneider et al. | https://imagej.nih.gov/ij/ |
| GraphPad Prism v8.0.1 | GraphPad | https://www.graphpad.com/ |
| Novoexpress v1.6.1 | Agilent Technologies | https://www.agilent.com/ |
| Biorender | Biorender | https://app.biorender.com/ |
| GEPIA2 | Tang, Z. et al. | http://gepia2.cancer-pku.cn/ |

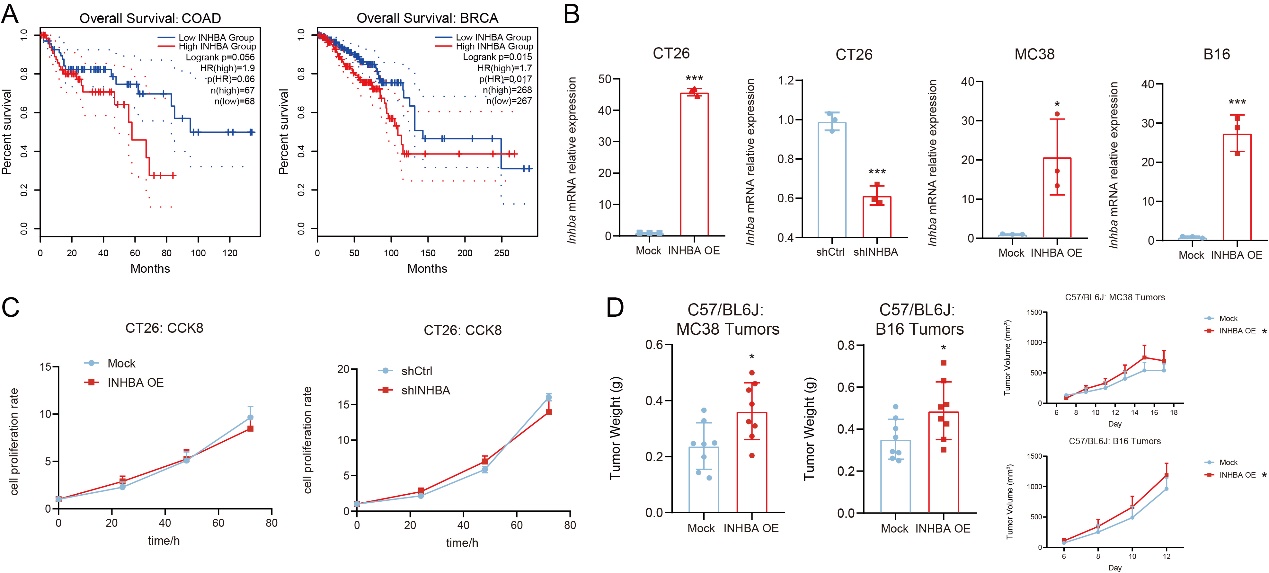

**FigS1. Effect of INHBA on tumor growth.** (**A**) The Kaplan-Meier curves show the overall survival in patients with colon adenocarcinoma (COAD) and breast cancer (BRCA) from the TCGA database determined by GEPIA. (**B**) Validation of INHBA gene-editing cells by qPCR (n = 3). (**C**) Cell proliferation assays (n = 6) were performed on CT26 INHBA gene-editing cells. (**D**) Tumor weights and growth kinetics of mock or INHBA OE MC38/B16 tumors in C57BL/6 mice (n = 8). The data are presented as the mean ± SEM. * p < 0.05; *** p < 0.001 by unpaired t test.

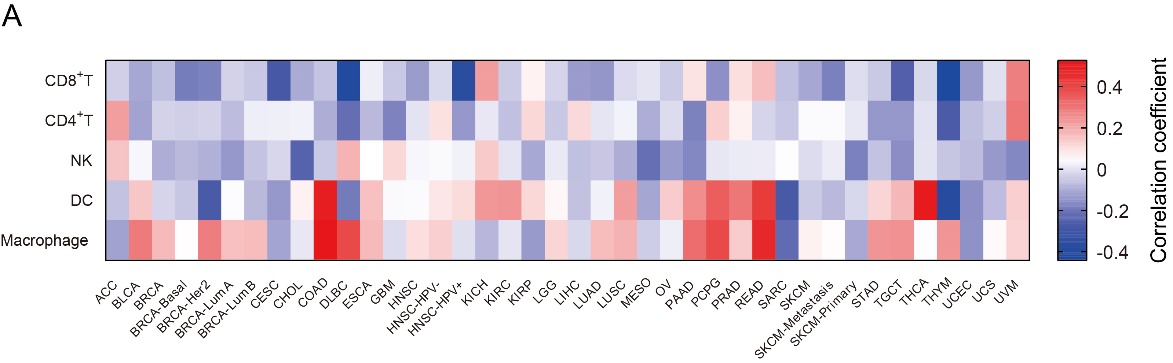

**FigS2. Correlation between tumor INHBA and immune cell infiltration.** (**A**) The heat map shows the correlation of *INHBA* expression with immune cell infiltration level in different tumor types as indicated in the TCGA database and was determined by TIMER2.0.

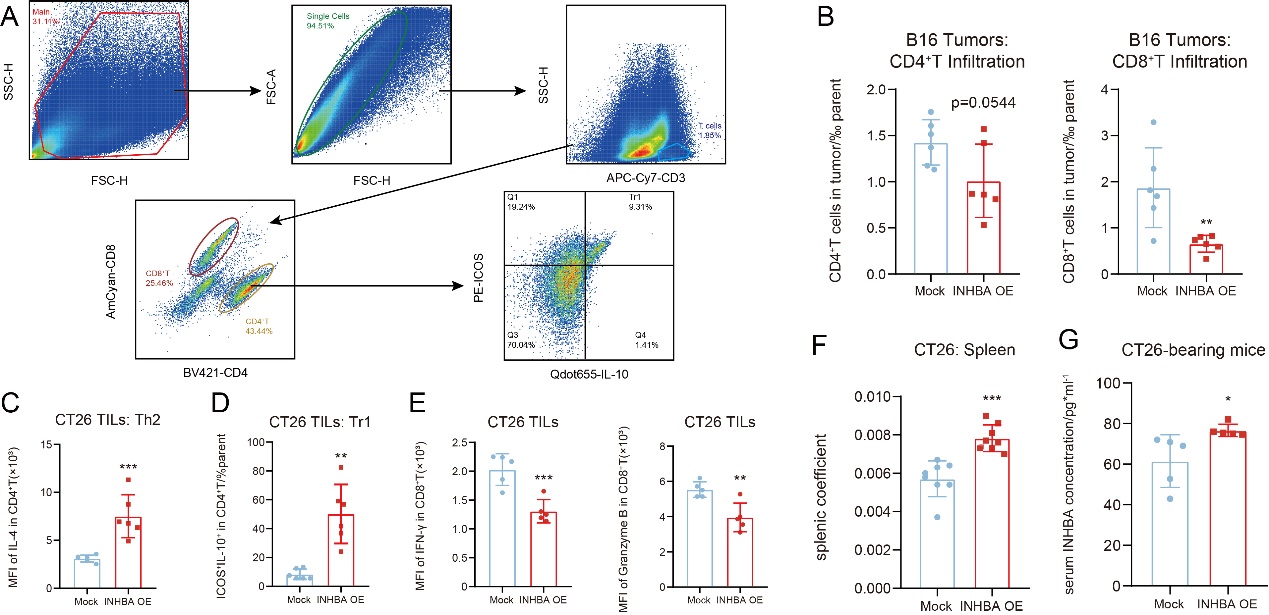

**FigS3. Effect of INHBA on tumor immunity.** (**A**) Representative flow-cytometry gating strategy for quantifying T cell subsets in murine tumors. (**B**) Flow cytometric analysis showing the percentage of CD4^+^T and CD8^+^T cells infiltration in mock or INHBA OE B16 tumors (n = 6). (**C)** Flow cytometric analysis showing the MFI of IL-4 in CD4^+^ T cells in mock or INHBA OE CT26 tumors (n = 6). (**D**) Flow cytometric analysis showing the percentage of Tr1 in CD4^+^ T cells in mock or INHBA OE CT26 tumors (n = 6). (**E**) Flow cytometric analysis showing the MFI of IFN-γ (left panel) and granzyme B (right panel) in CD4^+^ T cells in mock or INHBA OE CT26 tumors (n = 6). (**F**) Splenic coefficient of mock or INHBA OE CT26 tumor bearing mice was calculated (n = 8). Splenic coefficient equals spleen weight/body weight. (**G**) Concentration of INHBA in serum of mock or INHBA OE CT26 tumor bearing mice was determined by ELISA (n = 5). The data are presented as the mean ± SEM. * p < 0.05; ** p < 0.01; *** p < 0.001 by unpaired t test.

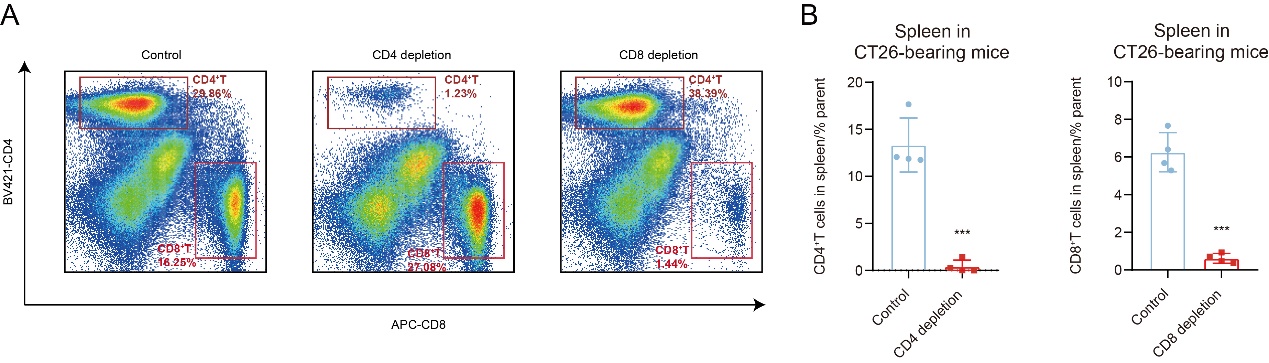

**FigS4. Validation of immune cell depletion efficacy.** (**A** to **B**) Representative flow-cytometry histogram (A) and quantification (B) for percentage of CD4^+^ T and CD8^+^ T cells in murine spleens with or without CD4^+^ T or CD8^+^ T cells depletion reagents (n = 4). The data are presented as the mean ± SEM. *** p < 0.001 by unpaired t test.

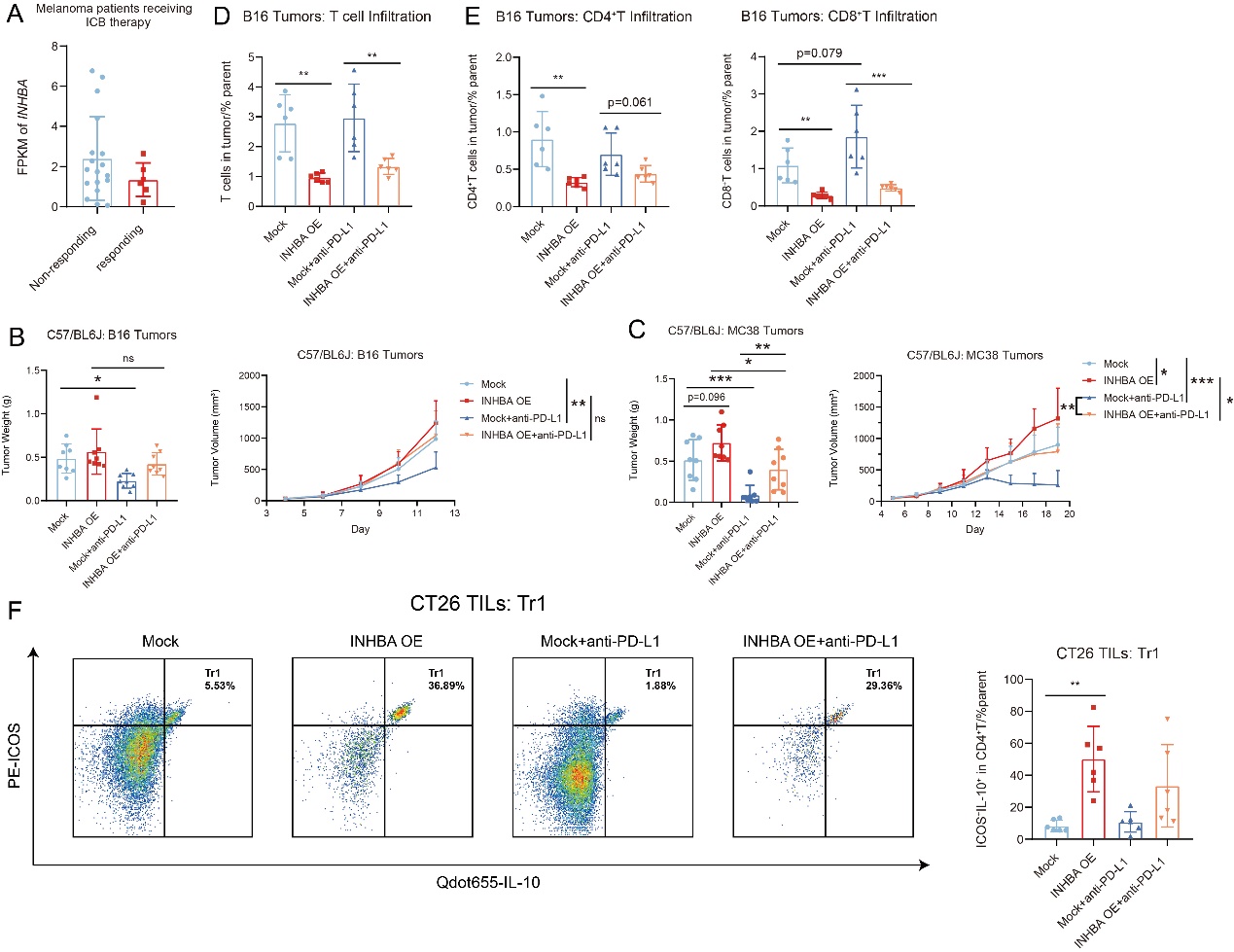

**FigS5. Effect of INHBA on anti-PD-L1 resistance.** (**A**) INHBA mRNA expression was compared between tumor from melanoma patients responsive or non-responsive to anti-PD-1/PD-L1 therapy provided by available GEO data (GSE91061). (**B**) Tumor weights (left panel) and growth kinetics (right panel) of mock or INHBA OE B16 tumors with treatment of atezolizumab or not (n = 8). (**C**) Tumor weights (left panel) and growth kinetics (right panel) of mock or INHBA OE MC38 tumors with treatment of atezolizumab or not (n = 8). (**D** to **E**) Quantification of T cells (D), CD4^+^ T and CD8^+^ T cells infiltration (E) from mock or INHBA OE B16 tumors with or without treatment of atezolizumab (n = 6). (**F**) Representative flow-cytometry histogram (left panel) and quantification (right panel) for percentage of Tr1 in CD4^+^ T cells from mock or INHBA OE CT26 tumors with or without treatment of atezolizumab (n = 6). The data are presented as the mean ± SEM. * p < 0.05; ** p < 0.01; *** p < 0.001; ns not significant by unpaired t test or one-way ANOVA followed by Tukey’s multiple comparisons test.

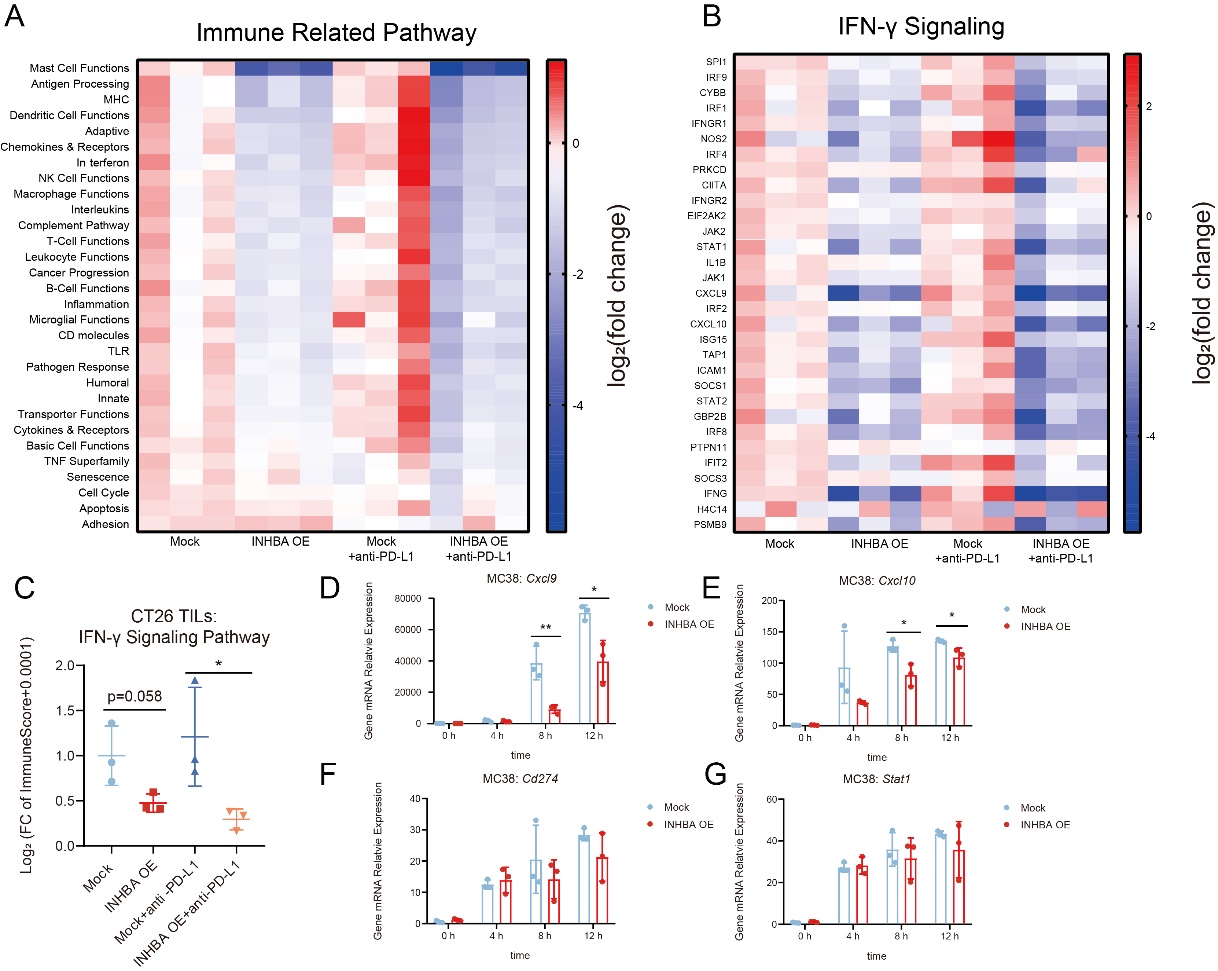

**FigS6. Effect of INHBA on IFN-γ response.** (**A**) Signature scores [defined as the mean log_2_ (fold change) among all genes measured by RNA-seq in the signature] for immune-associated processes are shown as heat map (n = 3). Samples are from TILs of CT26 bearing mice. (**B**) Expression scores [defined as log_2_ (fold change) of all genes measured by RNA-seq in the signature] for all genes in WP_TYPE_II_INTERFERON_SIGNALING_IFNG geneset are shown as heat map (n = 3). Samples are from TILs of CT26 bearing mice. (**C**) Signature scores for WP_TYPE_II_INTERFERON_SIGNALING_IFNG geneset are shown as scatter plot (n = 3). Samples are from TILs of CT26 bearing mice. (**D** to **G**) Gene expression for *Cxcl9* (D), *Cxcl10* (E), *Cd274* (F) and *Stat1* (G) in mock or INHBA OE MC38 cells with IFN-γ treatment (n = 3). The data are presented as the mean ± SEM. * p < 0.05; ** p < 0.01 by unpaired t test or one-way/two-way ANOVA followed by Tukey’s multiple comparisons test.

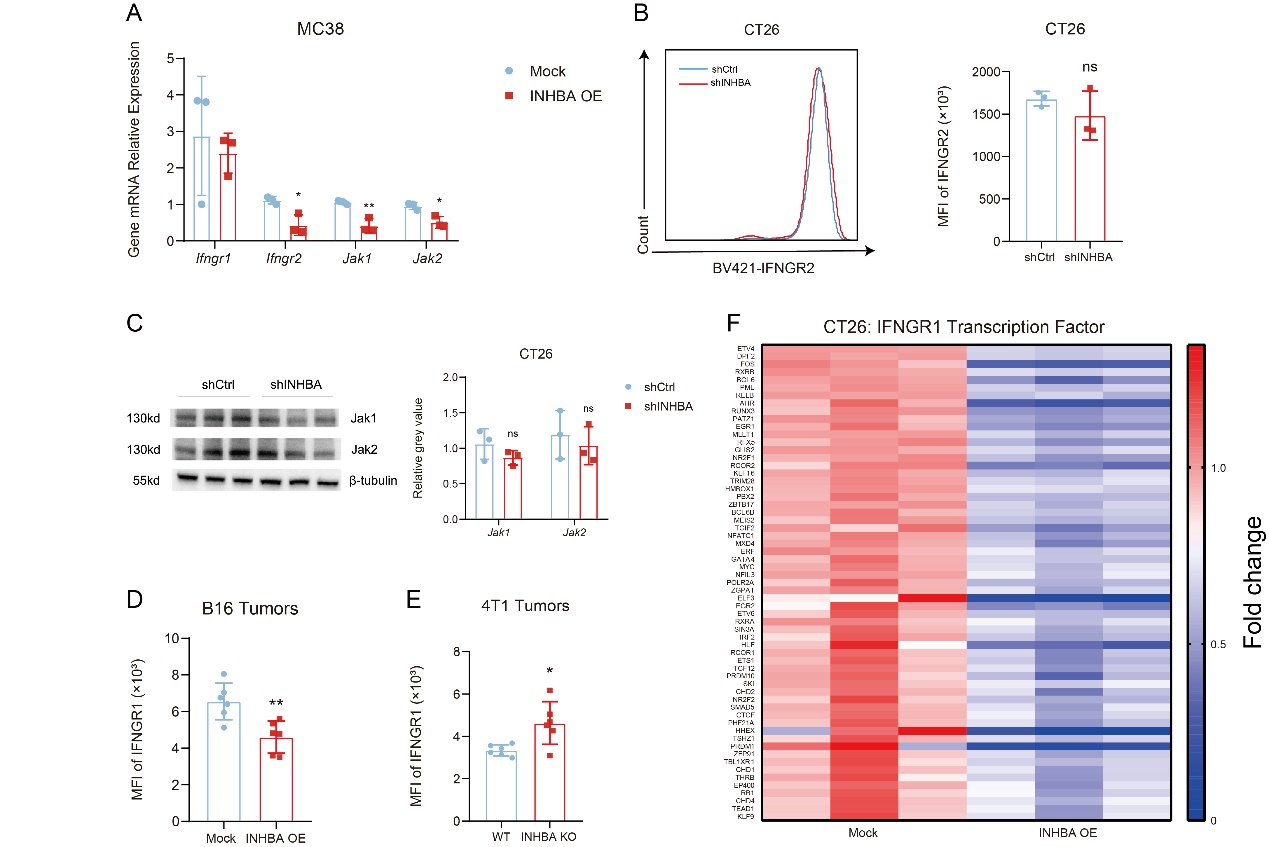

**FigS7. Effect of INHBA on expression of IFN-γ receptor complex.** (**A**) Gene expression for *Ifngr1*, *Ifngr2*, *Jak1* and *Jak2* in MC38 INHBA OE in comparison to MC38 mock cells (n = 3). (**B**) Representative histograms (left panel) and quantification analysis (right panel) showing the effect of *Inhba* knockdown on IFNGR2 expression in CT26 cells (n = 3). The level of IFNGR2 was detected by flow cytometry. (**C**) Western blot analysis of JAK1 and JAK2 expression in shCtrl or shINHBA CT26 cells (n = 3). β-Tubulin was used as the protein loading control. (**D** to **E**) Flow cytometric analysis showing the MFI of IFNGR1 in mock or INHBA OE B16 tumors (D; n = 6) and WT or INHBA KO 4T1 tumors (E; n = 6). (**F**) Expression scores [defined as fold change of genes measured by RNA-seq in the signature] for IFNGR1 transcription factors with significant difference between mock CT26 cells and INHBA OE CT26 cells are shown as heat map (n = 3). The data are presented as the mean ± SEM. * p < 0.05; ** p < 0.01; *** p < 0.001; ns not significant by unpaired t test.

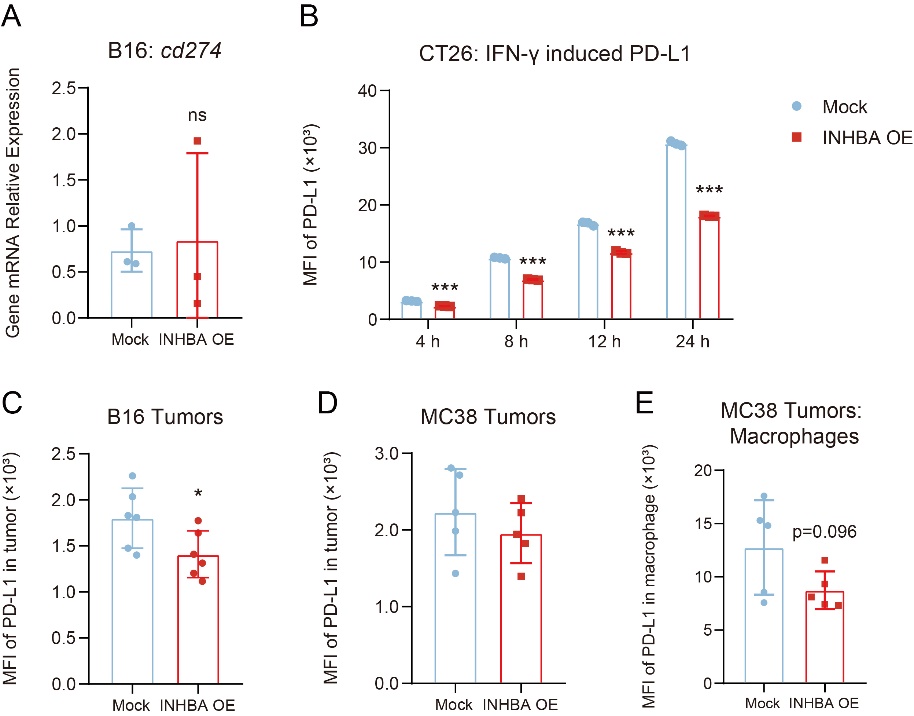

**FigS8. Effect of INHBA on tumor PD-L1 expression.** (**A**) Gene expression for *Cd274* in mock or INHBA OE B16 cells (n = 3). (**B**) Flow cytometric analysis showing the MFI of PD-L1 in mock or INHBA OE CT26 cells with treatment of IFN-γ for 4, 8, 12 and 24 h (n = 3). (**C** to **D**) Flow cytometric analysis showing the MFI of PD-L1 in mock or INHBA OE B16 tumors (C; n = 8) and MC38 mock or INHBA OE MC38 tumors (D; n=6). (**E**) Flow cytometric analysis showing the MFI of PD-L1 in macrophages of mock or INHBA OE MC38 tumors (n = 6). The data are presented as the mean ± SEM. * p < 0.05; *** p < 0.001; ns not significant by unpaired t test.

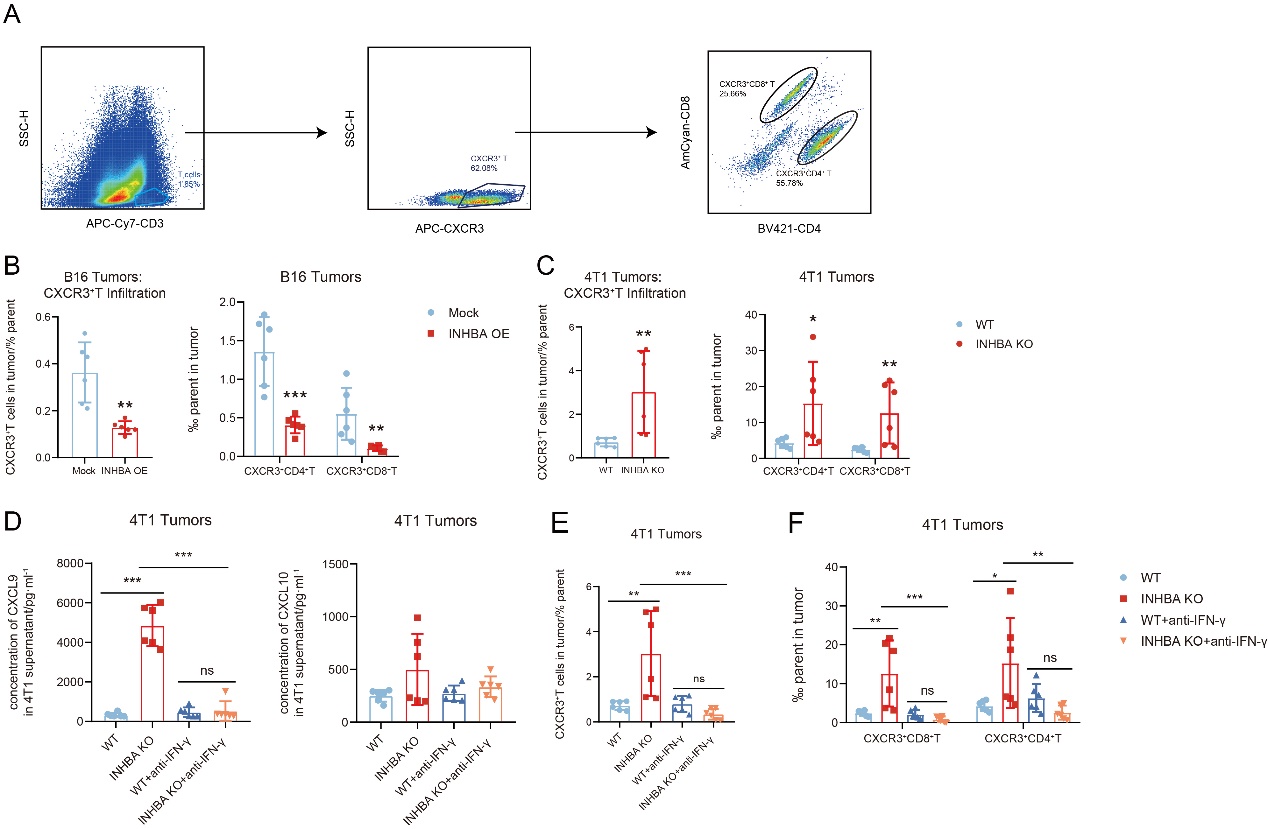

**FigS9. Effect of INHBA on CXCL9/CXCL10/CXCR3 axis.** (**A**) Representative flow-cytometry gating strategy for quantifying CXCR3^+^ T cell subsets in murine tumors. (**B** to **C**) Flow cytometric analysis showing the percentage of CXCR3^+^ T (left panel), CXCR3^+^ CD4^+^ T and CXCR3^+^ CD8^+^ T (right panel) cells infiltration in mock or INHBA OE B16 tumors (B; n = 6) and in WT or INHBA KO 4T1 tumors (C; n = 6). (**D**) The concentration of CXCL9 (left panel) and CXCL10 (right panel) were determined in WT or INHBA KO 4T1 tumors with or without treatment of anti-IFN-γ (n = 6). (**E** to **F**) Flow cytometric analysis showing the percentage of CXCR3^+^ T (E), CXCR3^+^ CD4^+^ T and CXCR3^+^ CD8^+^ T cells (F) infiltration in WT or INHBA KO 4T1 tumors with or without treatment of anti-IFN-γ (n = 6). The data are presented as the mean ± SEM. * p < 0.05; ** p < 0.01; *** p < 0.001; ns not significant by unpaired t test or one-way ANOVA followed by Tukey’s multiple comparisons test.

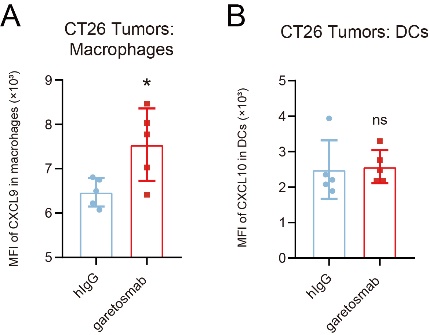

**FigS10. Effect of anti-activin A antibody on CXCL9 secretion from myeloid cells.** (**A** to **B**) Flow cytometric analysis showing the MFI of CXCL9 in macrophages (A) and DCs (B) of CT26 tumors treated with hIgG or garetosmab (n = 5). The data are presented as the mean ± SEM. * p < 0.05; ns not significant by unpaired t test.
